# This population does not exist: learning the distribution of evolutionary histories with generative adversarial networks

**DOI:** 10.1101/2022.09.17.508145

**Authors:** William W. Booker, Dylan D. Ray, Daniel R. Schrider

## Abstract

Numerous studies over the last decade have demonstrated the utility of machine learning methods when applied to population genetic tasks. More recent studies show the potential of deep learning methods in particular, which allow researchers to approach problems without making prior assumptions about how the data should be summarized or manipulated, instead learning their own internal representation of the data in an attempt to maximize inferential accuracy. One type of deep neural network, called Generative Adversarial Networks (GANs), can even be used to generate new data, and this approach has been used to create individual artificial human genomes free from privacy concerns. In this study, we further explore the application of GANs in population genetics by designing and training a network to learn the statistical distribution of population genetic alignments (i.e. data sets consisting of sequences from an entire population sample) under several diverse evolutionary histories—the first GAN capable of performing this task. After testing multiple different neural network architectures, we report the results of a fully differentiable Deep-Convolutional Wasserstein GAN with gradient penalty that is capable of generating artificial examples of population genetic alignments that successfully mimic key aspects of the training data, including the site frequency spectrum, differentiation between populations, and patterns of linkage disequilibrium. We demonstrate consistent training success across various evolutionary models, including models of panmictic and subdivided populations, populations at equilibrium and experiencing changes in size, and populations experiencing either no selection or positive selection of various strengths, all without the need for extensive hyperparameter tuning. Overall, our findings highlight the ability of GANs to learn and mimic population genetic data and suggest future areas where this work can be applied in population genetics research that we discuss herein.

## INTRODUCTION

Accurately capturing the properties of genetic variation to infer evolutionary processes is a major goal of population genetics. Throughout the history of population genetic research, myriad summary statistics have been created to accomplish this task. Ideally, the values of a given statistic follow theoretical predictions and are robust to additional unconsidered forces. For example, *π* (Nei and Li 1979), which measures the degree of nucleotide diversity in a population, is reduced by linked directional selection in a manner that is predicted by the selection coefficient and the recombination landscape around the selected site(s) (Smith and Haigh 1974; Kaplan *et al*. 1989; Hudson and Kaplan 1995; Nordborg *et al*. 1996). However, diversity levels are also affected by other processes such as population bottlenecks and population subdivision (Simonsen *et al*. 1995). Although the simplicity of this univariate summary of genetic diversity makes it well-suited for theoretical studies of the effects of various evolutionary processes on diversity, this also renders *π* relatively uninformative for distinguishing among the diversity of complex evolutionary scenarios considered in modern population genetic inference.

As a solution to this problem, researchers have turned to either creating more complex statistics (Voight *et al*. 2006; e.g. Ferrer-Admetlla *et al*. 2014) or using methods that rely on a number of theoretically informative summary statistics to infer a process or evolutionary history (Beaumont 2010; Schrider and Kern 2018). Common to the latter approaches is the leveraging of increasingly greater amounts of information or observations to make evolutionary inferences, and this increased inferential power typically comes at the cost of reduced interpretability. As the number of summary statistics used increases, it becomes less apparent how any one statistic, and especially any combination of statistics, is informative about the process in question. Ultimately, the tools we use to make meaningful inference currently lie on a continuum between interpretability and power, and researchers must make a choice within this continuum for the question at hand.

If simple statistics lie on one end of the interpretability-power continuum, deep-learning methods lie on its opposite. Often described as a ‘black box’, deep-learning models frequently have millions of parameters that make identifying the features responsible for a given inference a challenging task (although research is being conducted on this task, see: Simonyan et al. 2013; Selvaraju et al. 2017; Burkart and Huber 2021). However, these methods can be remarkably powerful and in some cases have made dramatic progress in the biological domain (e.g. Jumper *et al*. 2021). In population genetics, deep-learning methods have been used for a number of purposes, such as detecting introgression (Flagel *et al*. 2019)—including adaptive introgression (Gower *et al*. 2021); distinguishing among various modes of natural selection as well as neutral evolution (Schrider and Kern 2016; Kern and Schrider 2018; Isildak *et al*. 2021; Whitehouse and Schrider 2022); estimating parameters such as selection coefficients and recombination rates (Flagel *et al*. 2019; Adrion *et al*. 2020a), dispersal distances (Smith *et al*. 2022), and population sizes (Sanchez *et al*. 2021); and visualizing population structure (Battey *et al*. 2021).

Although deep-learning networks are often thought of as an uninterpretable ‘black box’, one argument suggests that such algorithmically generated models may in fact be well suited to model natural phenomena that are generated by complex processes that are inherently uninterpretable through the lens of simple statistical models (Breiman 2001). Rather than reducing our data down to some predetermined set of variables, many of these networks—such as those that use convolutional neural networks (LeCun et al. 1989; LeCun et al. 1998)—are able to learn their own data representations from the raw data itself. Because they do not rely on *a priori* assumptions of how to summarize our data, these methods can potentially learn to extract the information needed to make accurate inferences without requiring our data be manipulated in ways that may be biased or otherwise omit useful information. While much of the natural world can be understood using mechanistic models with clear causes and effects, restricting ourselves to this frame of understanding renders biological science myopic— precluding our ability to discover more complex phenomena with emergent properties otherwise unaccounted for. In evolutionary biology, and in particular population genetics, progress has traditionally relied on this mechanistic framework through the development of theoretical models that are then used to explain our observations (Schrider and Kern 2018). Deep-learning methods, however, allow us to take the opposite approach: letting observation directly inform our understanding (LeCun *et al*. 2015).

One limitation of this approach is that for most evolutionary processes we have few example phenomena where the true evolutionary history with respect to the phenomenon of interest is known with complete certainty. For example, although there are many loci where we strongly suspect there has been recent positive selection, even for the best studied examples we often cannot be certain of the identity of the selected site, or even gene, let alone of parameters such as the strength of selection. Thus, typically in machine learning applications to population genetic problems, the training observations themselves must be generated from simulations based on reductionist models (but see Schrider and Kern 2015). However, if we leverage the usefully reductive aspects of biology (e.g. predictable changes in allele frequencies in response to selection) alongside the advantages of deep-learning methods to factor in the unreducible parts (i.e. the messy nature of biological data and its collection), it may be possible to render this limitation advantageous.

In this study, we focus on Generative Adversarial Networks (GANs), a specific class of deep-learning network that may be particularly useful in population genetics. Variations of GANs have been previously used to generate artificial human genomes (Yelmen *et al*. 2021) and estimate demographic parameters of populations (Wang *et al*. 2020). While the explicit function of these networks is to generate artificial examples of some set of input data, training these networks aims to learn the statistical distribution of the input data in a multidimensional parameter space (Creswell *et al*. 2018). As a result, GANs can be useful for any task where identifying this distribution is advantageous and can be exploited, such as data augmentation (Bousmalis *et al*. 2017), classification (Radford *et al*. 2015), clustering (Kim and Ha 2021), or the transfer of properties from one dataset onto another (i.e. style transfer; Karras *et al*. 2018). Importantly, by learning this distribution, any artificial examples generated by the GAN are not tied to any one input example. This property makes GANs particularly useful for generating testing data containing sensitive information or where privacy is a concern, such as data collected from human subjects (Yale *et al*. 2019; Yelmen *et al*. 2021).

Here, we complement previous work using generative models in population genetics by designing the first fully differentiable GAN capable of generating artificial population genetic alignments. Previous work has shown that GANs can learn to generate individual genomes from a shared ancestry (Yelmen *et al*. 2021), analogous to drawing multiple individuals from a single ancestral recombination graph (ARG). Here, we demonstrate that GANs can simultaneously generate entire samples of genomes from the full distribution of samples that may be produced under a given evolutionary model, analogous to drawing an entire ARG from the distribution of ARGs produced by an evolutionary process. While qualitatively different in the data learned and generated, the architectural differences in our GAN may also allow us to capture additional information about the joint history of individuals in each sample. We begin with a design based on the Wasserstein GAN (Arjovsky *et al*. 2017) and test several different architectures to find the optimal design for this task. We then simulate alignments under a variety of evolutionary scenarios, including models of population size change, subdivision, and natural selection, and evaluate our GAN’s ability to accurately capture and replicate alignment properties under each of these scenarios. Finally, we discuss the strengths and weaknesses of our GAN, and propose paths forward for using GANs for population genetic modeling and inference.

## METHODS

### GAN Overview

Generative Adversarial Networks (GANs) are in actuality a combination of two different networks that compete against each other during training. As initially designed, these two networks consist of a generator that produces some output from a latent noise vector as input, and a discriminator that takes in both real and generated data and attempts to distinguish between the two (Goodfellow *et al*. 2014). By training these two networks in tandem, the generator should get progressively better at producing realistic copies while the discriminator gets better at distinguishing those fake copies from real ones. Critically, the generator is blind to the data it is trying to replicate, and only receives updates from the discriminator on its performance.

### Architecture and Implementation

All networks were created using python version 3.9.7 along with pytorch version 1.9.1 (Paszke *et al*. 2019) and numpy version 1.20.3 (Harris *et al*. 2020). To design our GAN, we tested several network architectures with different combinations of layers and properties to find an architecture that performed consistently across runs and population genetic models. Largely, the networks tested were variations of a Deep Convolutional GAN (Radford *et al*. 2015) or a ResNet GAN (as implemented in Gulrajani *et al*. 2017), with the ResNet GAN being largely a modification of the DCGAN architecture where the input to each convolutional layer block is added to its output. These networks were initially designed to reproduce two-dimensional images with color values on 3 channels, and this data structure is easily translatable to population genetic alignments with biallelic polymorphisms by reducing the network to a single color channel, analogous to a black and white image, with values that represent ancestral (0) or derived (1) alleles. However, while this structure captures allelic information at each site it leaves out valuable information about the position of those sites along a sequence. To include this information, in both the generator and discriminator networks we included two branches, one performing two-dimensional convolutions on the alignment matrix, and the other performing one-dimensional convolutions on the vector of positions for each polymorphism in the alignment (Fig. 1).

**Figure 1.**
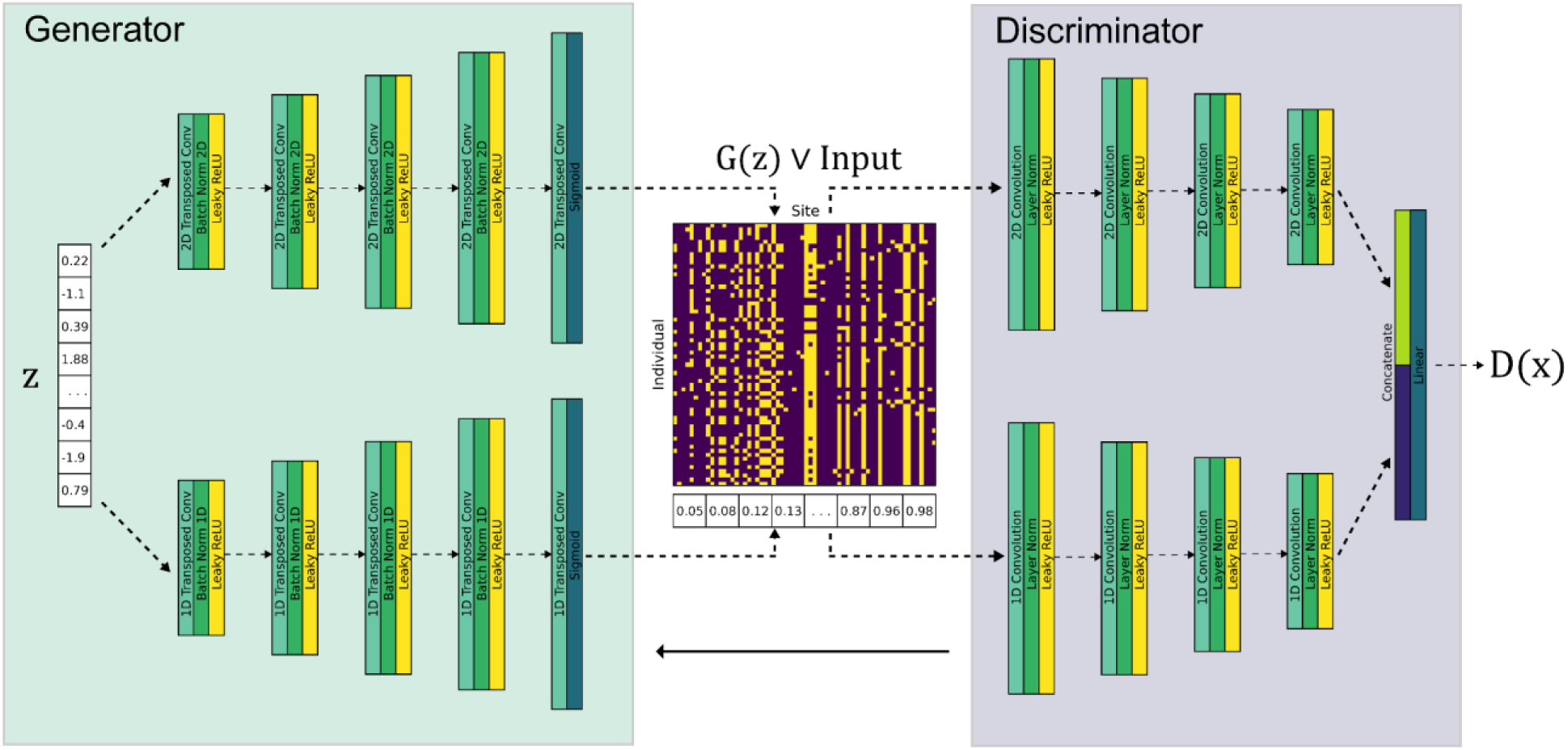
Architecture and implementation of our Generative Adversarial Network. The final network chosen is a branched Deep Convolutional Gan (DCGAN) with Wasserstein loss (WGAN) and a gradient penalty to enforce 1-Lipschitz continuity. The generator takes in a random noise vector drawn from a multidimensional Gaussian and is branched into independent transposed convolutional layers to produce artificial alignments along with a vector of the positions of segregating sites. The discriminator takes in either generated or real alignments and positions vectors and evaluates them through a branched network of convolutions. Loss is calculated using the earth mover’s distance between real and generated alignments with an additional gradient penalty term, and is used to train the discriminator which in turn trains the generator.

One difficulty with GANs as initially designed is the “vanishing gradient” problem, where the discriminator learns too quickly and the generator is unable to keep up. To prevent this from occurring, we used the Earth Mover’s (EM) distance as our loss function as implemented in the Wasserstein GAN (WGAN; Arjovsky *et al*. 2017). Rather than treating the discriminator and generator as adversaries in a min-max game using the discrete prediction of real (1) or fake (0) as in the original GAN (Goodfellow et al. 2014), the WGAN attempts to minimize the ‘effort’ required to transform the distribution of values output by the discriminator (or critic, in Wasserstein GANs) from generated examples into the distribution of values output by real examples. However, by using the EM distance as a loss function the network is no longer 1-Lipschitz continuous and requires either weight clipping or other modifications to enforce the Lipschitz constraint. To enforce this constraint, we added a gradient penalty (Gulrajani *et al*. 2017) to the loss calculation:

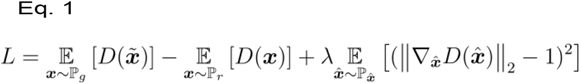

Here, the first two elements of the equation 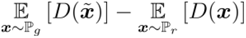 calculate the EM distance between the discriminator’s predictions on generated (the first term) and real data (the second term), with the second half 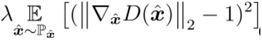 calculating a penalty on the gradient norm of the discriminator for examples randomly sampled in between real and generated data 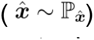, with deviations of this norm from 1 being penalized according to the penalty hyperparameter λ. By designing our GAN this way, training should converge more easily and be more stable across runs and hyperparameter values. Finally, we used the ‘Adam’ optimizer for training (Kingma and Ba 2014).

Within the overall architectures outlined above, we also tested a number of more subtle variations of the network architecture to arrive at a more optimal solution. These included various normalization layers following convolutions (batch, instance, layer), activation functions following internal convolutions (ReLU and leaky ReLU), adding Gaussian noise between layers in the discriminator, upsampling and transposed convolutions in the generator, and final activation layers in the generator (sigmoid, tanh, Gumbel softmax). Regarding the final activation, the sigmoid activation [0,1] value was interpreted as the probability of having a derived allele for a given individual at each site, with the tanh activation [-1,1] being a similar probability when re-scaled to [0,1]. The Gumbel softmax activation was tested in an attempt to discretize the generated alignments fed into the discriminator with the value for a given individual at each site representing an allele belonging to either the ancestral (0) or derived (1) class.

In addition to these variations, we also attempted to add in additional information in the models related to the variation and frequency of alleles. A fundamental difference between simulated alignments and alignments generated by a GAN is that output from simulated alignments are guaranteed to have variation at every site, whereas the GAN has no such guarantee and variation at every site must be learned or achieved by some other means after generating the data if generating alignments with the same number of sites as the input is desired. Although alignments don’t need to have the same number of sites for comparison, early tests suggested not requiring variation led to a reduction in singleton sites almost exclusively (Supp. Fig. 1).

To assist training in ensuring variation at every site, we tested networks that had an additional loss term penalizing monomorphic sites added to the generator loss (Eq. 2, where *i* is the number of sites and ω is some weight either fixed or tied to the gradient penalty loss) and additional features passed as input into the discriminator including the sum, minimum, maximum, and variance of values at each site. To ensure variation at every site in generated alignments, a random individual at sites initially monomorphic for the ancestral allele was chosen to have the derived allele, with this choice weighted by each individual’s probabilities from the final activation layer. The same was done for sites monomorphic for the derived allele, except in these cases the weight used to randomly select an individual to be given the ancestral allele was one minus the probability from the final activation later.

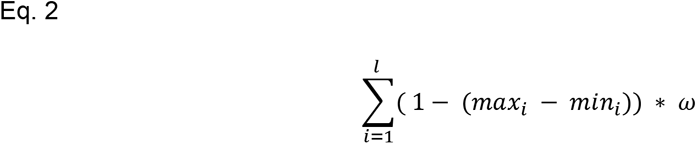

### Simulations and Models

We tested the ability of our GAN to broadly capture the properties of population genetic alignments by training the GAN using several different simulated models as input. All models were simulated using coalescent simulators: ms (Hudson 2002), discoal (Kern and Schrider 2016), or stdpopsim version 0.1.2 (Adrion *et al*. 2020b) with msprime version 1.0.2 (Kelleher *et al*. 2016; Baumdicker *et al*. 2022) as the backend of stdpopsim. The most basic of models assessed were a standard neutral model both with and without recombination. Together, these models allow us to evaluate the GAN’s ability in the simplest of cases as well as its ability to capture information about the covariance of nearby sites when recombination is present. We also evaluated a model of isolation with migration of two equally sized populations, experimenting with several rates of migration. To evaluate the GAN’s ability to learn to model population structure from structure in the alignment, we trained the GAN where the input was the direct output from the simulation (where the population 1 is the first 32 rows of the alignment, and population 2 the remaining 32 rows) and where the input rows (individuals) were shuffled. Additionally, we evaluated models where there was a recent population expansion or contraction. To assess performance on a more complex demographic model, we trained the GAN using simulations from the two-population ‘Out of Africa’ model from Tennessen et al. (2012).Finally, we assessed the ability of the GAN to replicate patterns of recent positive selection by simulating models of strong and moderate selective sweeps (2*Ns*=2000 and 2*Ns*=100, respectively) where the selected site was in the middle of the alignment and the sweep had completed upon sampling. The standard neutral and IM models were simulated with a set number of segregating sites using the ms ‘-s’ flag. All other models were simulated with a variable number of sites, and 64 sites were extracted for training by taking the 64 sites surrounding the most central position in the alignment. Note that because of this, the locus-wide recombination rate parameter used for these simulations is greater than or equal to that of the window of the alignment fed as input into the GAN. This also ensured that, for the simulations of selection, the selected site was always in the center of the alignment and could be properly captured by windowed diversity metrics. All simulation commands are available in the supplement (Supp. Table 1).

### Evaluation

We used a number of measures relevant to population genetics to evaluate the performance and training of our GAN. Because of the nature of adversarial training, the loss of GANs does not directly correlate to the ability of the GAN to generate realistic copies. Typically, these GANs are evaluated by some metric calculated using a pretrained network to determine the similarity of GAN output to the input data, such as inception scores (Salimans *et al*. 2016), however the inception network was designed and trained for use with 3-channel color images rather than the binary alignments used here. To evaluate our single channel output, we evaluated our GAN using the 2D sliced Wasserstein distance of the distribution characterized by the first two principal components calculated from the site-frequency spectra (SFS) of the input and generated alignments (referred hereafter as 2DSWD). This measurement is an approximation of the exact Wasserstein (or earth mover’s) distance calculated through linear slicing of the two distributions (Bonneel *et al*. 2015).

Similar to Yelmen et al. (2021), we used nearest-neighbor adversarial accuracy (here calculated using the SFS of each alignment) to evaluate over and underfitting (see Yale *et al*. 2019). We used Eq. 3-5 (reproduced from Yelmen et al. (2021)) to calculate 3 adversarial accuracy scores: AA_truth_, AA_syn_, and AA_TS_. Here, *m* is the number of alignments (real alignments for Eq. 3, and generated alignments for Eq. 4), the function 1 returns 1 if its input value is true and 0 otherwise, d_TS_(i) is the distance between the ith real alignment its nearest generated alignment, d_ST_(i) is the distance between the ith generated alignment and its nearest real alignment, d_TT_(i) is the distance between the ith real alignment indexed and its nearest neighboring real alignment, and d_SS_(i) is the distance between the ith generated alignment and its nearest neighboring generated alignment. The optimal value for all AA values is 0.5, with an AA_TS_ above and below 0.5 suggesting the generator is underfitting or overfitting, respectively.

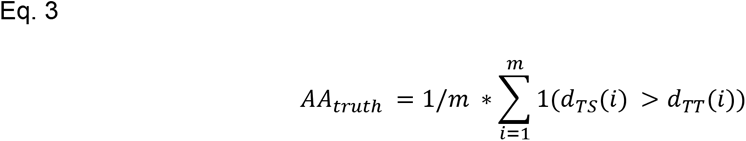

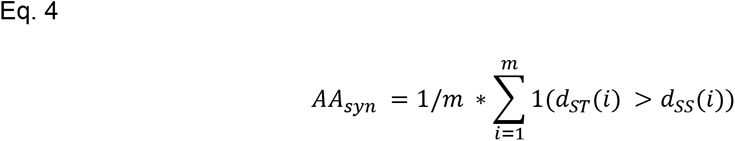

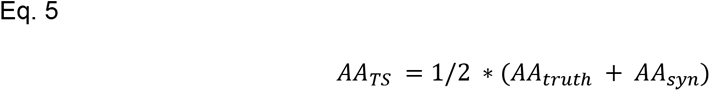

To broadly capture the similarity of the input and generated alignments, we calculated and compared the average SFS, and joint SFS for two-population models using scikit-allel (Miles *et al*. 2021). To assess the GAN’s ability to mimic recombination, we calculated the decay of gametic linkage disequilibrium (as measured by *r*^2^) with distance. To investigate population size histories consistent with the observed SFS, we used the program Stairway Plot version 2.1.1 (Liu and Fu 2020). Finally, we also calculated the summary statistics Watterson’s *θ* (Watterson 1975), Tajima’s *D* (Tajima 1989), *π* (Nei and Li 1979), and *ω* (Kim and Nielsen 2004). For each model evaluated, we used a training set of 20,000 input alignments and trained for 10,000 epochs. Because GANs may become unstable as training goes on, we saved the network after every 100 epochs and evaluated the performance of the saved model that minimized the 2DSWD as calculated above.

## RESULTS

### Architecture

#### Final architecture

We experimented with several GAN architectures for generating population genetic alignments (Methods). Of the architectures tested, our final design was a modification of the DCGAN with Wasserstein loss and gradient penalty to enforce 1-Lipschitz continuity (Fig. 1). The final design takes in a noise vector of samples from a 128-dimensional standard normal distribution as input. This input is then branched into two series of convolutions to produce the alignment and positions vector, respectively. In the generator, the initial transformation from the latent vector in the alignment branch involves a 2D transposed convolution from the 128-channel noise vector to 512 channels with a kernel size of 4, a stride of 1, and no padding, followed by a batch normalization layer and leaky ReLU activation with a negative slope of 0.1. Following the initial transformation are 3 additional transposed convolutional layers with batch normalization and leaky ReLU, and each of these layers reduces the number of channels by half with a kernel size of 4, stride of 2, and padding of 1. The final transposed convolution has the same kernel, stride, and padding as previous layers and produces a 64×64 alignment on a single channel, has no batch normalization layer, and is followed by a sigmoid activation layer producing values from 0 to 1 which we interpret as the probability of having a derived allele for each individual at each site. For the positions branch, the same 128-channel noise vector used in the alignment branch is passed through an initial 1D transposed convolution that reduces the number of channels by half with a kernel size of 4, stride of 1, and no padding, followed by a batch normalization layer and a leaky ReLU activation with negative slope 0.1. This initial transformation is followed by 3 additional 1D transposed convolutional layers that each reduce the number of channels in half and are each followed by batch normalization and leaky ReLU activation, and that have a kernel size of 4, stride of 2, and padding of 1. The final 1D convolution produces a 1×64 vector on a single channel with a single convolution and is followed by a sigmoid activation layer to produce values from 0 to 1, with the boundaries of this range indicating the beginning and end of the chromosome, respectively.

The discriminator largely mirrors the generator in the reverse. In the alignment branch, the 64×64×1 alignment is initially transformed into 64 channels via a 2D convolution with a kernel size of 4, stride of 2, and padding of 1, followed by a layer normalization layer and a leaky ReLU activation with negative slope 0.1. This is followed by 3 additional 2D convolutions with the same layer normalization and leaky ReLU activation, but instead the number of channels doubles after each of these convolution layers. The positions branch is initially transformed through a 1D convolution into 16 channels with a kernel size of 4, stride of 1, and no padding, followed by layer normalization and leaky ReLU activation with negative slope 0.1. This initial transformation is then followed by 3 additional 1D convolutions with the same layer normalization and leaky ReLU activation, but with the number of output channels doubling after every convolution. After all convolutions, the alignment and positions branches are both flattened down to a single dimension, concatenated, and passed through a dense layer with a linear activation to produce a single output.

Hyperparameter tuning produced some variation in the performance of our GAN, but largely the GAN demonstrated stability over a wide variety of hyperparameter values (Supp. Table 2). Additionally, there was no clear value for any hyperparameter that consistently performed better than others across multiple hyperparameter combinations. Ultimately, we chose the hyperparameter values that minimized the average of the last 5 saved 2DSWDs. This distance was calculated and saved after every 100 epochs, training for 5000 epochs total with 10,000 alignments as input. The final hyperparameter values were: batch size = 256, lambda = 10, critic iterations = 5, negative slope = 0.1, and learning rate = 0.0002.

#### Additional architectures

Aside from our final GAN design, we tested several modifications of the final architecture or alternatives that we found performed more poorly. Most surprisingly, ResNet architectures (He et al. 2015; Arjovsky et al. 2017), which show improved performance over DCGANs in generating color images, performed worse than DCGANs for our purposes. Rarely did ResNet architectures produce something resembling an alignment, and when it did, these alignments were poor representations of the input alignments. Similarly, other proposed additions or variations to improve performance including upsampling in the generator, adding Gaussian noise between discriminator layers, smoothing allele values (subtracting or adding a small noise value drawn from U(0,0.05) to all values in the alignment), and flipping (switching 1-5% of generated and input example labels going into the discriminator) also reduced performance. Upsampling in particular produced output that had large, smooth blocks of similar values instead of the rigid, blocked structure of population genetic alignments. Using normalization layers other than batch normalization and layer normalization in the generator and discriminator, respectively, severely reduced performance–likely in part as a result of the Wasserstein loss function used (Arjovsky et al. 2017).

In addition to variations to the network structure, we also tried modifications to the architecture that introduced some penalty or evaluation to coerce the GAN to produce output that was more ‘alignment-like’. To attempt to ensure our alignments had variation at every site, we added a penalty term to the loss function weighted by one minus the difference between the minimum and maximum values at each site multiplied by some fraction of the gradient penalty (as the gradient penalty and loss magnitude increases during training). However, this penalty was either too weak to be effective, or when effective, produced alignments with site frequencies that avoided singletons or nearly fixed derived alleles (Supp. Fig. 2).

Because the added penalty resulted in poor behavior during training, we tried modifications that were implemented within the GAN itself to achieve variation at every site. Here, we added tensor operations into the features concatenated before the discriminator’s final linear activation, including the minimum, maximum, mean, and variance at every site. We investigated the effects of different combinations of these operations, upsampling the operations through a Multi-Layer Perceptron (MLP), and downsampling the other features through a dense layer. While these additions did not substantially harm performance, none produced better alignments than the DCGAN without these extra features.

As the input alignments have values of only 0 and 1, while the generated alignments can produce any value from 0 to 1 at any individual and site, we considered the possibility that this difference between discrete and continuous values negatively affected training. One possible fix for this that is fully differentiable is to have the generator produce log probabilities of belonging to one of two classes at each individual and site (i.e. ancestral and derived) instead of ending with a sigmoid activation, and to discretize the class affinity by sampling from the Gumbel-Softmax distribution (Jang et al. 2016; Maddison et al. 2016). While this Gumbel-Softmax variation did not dramatically reduce performance, we obtained better alignments when using the sigmoid-activated DCGAN, regardless of whether the former included the now-discretized tensor operations (i.e. the minimum, maximum, mean, and variance at every site) as input features for the discriminator as described above.

### Performance

In general, our GAN achieved the most drastic increases in similarity between input and generated alignments early in training (Supp. Fig. 3). For most models, the minimum 2DSWD was reached early in training (median of 4000 epochs), and continual training generally resulted in a gradual decrease in similarity, albeit with significant fluctuations, over the rest of training. However, some models, such as the population contraction and subdivided population with a migration rate of 0.1, remained relatively stable over training. For both selection models, there is an overall increase in 2DSWD after early training epochs, but with several later sharp decreases in 2DSWD that result in values well below the early training minimum. The shuffled migration model had the most unique training trajectory, exhibiting an overall decrease in 2DSWD over time, but with much larger fluctuations across epochs, as well as much higher 2DSWD values, than observed for other models.

We also assessed our models’ tendency to over or underfit the training data using adversarial accuracy measures (Methods; Yelmen et al. (2021)), summarizing each simulated or generated alignment by its SFS. For most models, AA_TS_ approached intermediate levels between 0.6 and 0.8 early in training and remained stable (Supp. Fig. 3). No model reached an AA_TS_ below 0.5 at any time during training, indicating a tendency of our GAN to underfit, rather than overfit, to the models (2021). Additionally, AA_truth_ values were generally close to 0.5 indicating that our simulated input alignments were on average roughly as similar to generated alignments as they are to other input alignments. AA_syn_ values, however, were generally closer to 1.0, indicating most generated alignments were closer to other generated alignments than the input alignments. In general terms, this trend indicates that the distribution of generated alignments overlapped the majority of the distribution of input alignments, but that a substantial fraction of the generated alignments were outside of the input alignment distribution and/or were clustered within regions of the distribution that had a comparatively low density of input alignments.

However, a few models show different adversarial accuracy trends than those of the general trend. For the subdivided population model with a migration rate of 0.1, both shuffled and unshuffled, all AA values remained tightly linked throughout most of the training run, diverging slightly after ∼4000 epochs. For the two population Out-of-Africa and moderate selection models, although the AA_TS_ value slowly increased after reaching a minimum close to 0.5, all AA values became more tightly linked as training went on. Finally, the strong selection model was unique in that the AA_syn_ scores were substantially lower than AA_truth_ scores throughout the training run, with the former closer to 0.5 and the latter closer to 1.0, while for most models we observed the opposite pattern. This pattern would indicate that the GAN was accurate in capturing a subset of the distribution of sweep alignments with strong selection, but not the whole distribution, and/or that the simulated alignments are found at relatively high density in some regions of the distribution where the generated data were at low density. We emphasize that these adversarial accuracy measures were based on the SFS, and thus it is possible that somewhat different patterns would emerge from an examination of other summaries of the alignment data.

### GANs can generate data mimicking a variety of population genetic models

#### Standard neutral model

To understand how our GAN performs in capturing the properties of the most basic models, we first tested its performance generating alignments simulated under a standard neutral model (i.e. a single panmictic population with no population size changes or natural selection). Overall our GAN was able to generate alignments under our most basic neutral model that broadly have the same properties and summary statistics of the input alignments. To visualize the similarity of simulated and generated alignments, we performed principal component analysis (PCA) of the site frequency spectra of these alignments. The first two principal components, shown in Fig. 2a, demonstrate that the generated alignments largely fall within the same distribution as the input alignments, although the generated alignments are more densely packed within a subregion of this distribution and thus do not capture the full distribution of input alignments. Although the bulk of the distribution is captured, GANs can also suffer from mode collapse (see Thanh-Tung and Tran 2020), which may be happening here as indicated by the elevated AA_syn_ values (Supp. Fig. 4). The SFS averaged across 1000 alignments (Fig. 2b), the distribution of positions of polymorphisms in the alignment (Fig. 2c), and nucleotide diversity distribution (Fig. 2d) are highly similar between the simulated and generated alignments. To test the GAN’s ability to implicitly model recombination, we added a recombination parameter to our standard neutral simulation. The amount of LD was lower in both the non-recombinant (Fig. 2e) and recombinant (Fig. 2f) neutral models’ generated alignments relative to the corresponding simulated alignments, likely due to noise in the generated alignments; the slight excess of singleton polymorphisms in the generated alignments is consistent with this explanation. Although the magnitude of LD was somewhat diminished, our GAN clearly produces the expected decay of LD with increasing distance between polymorphisms when recombination is added to the model.

**Figure 2.**
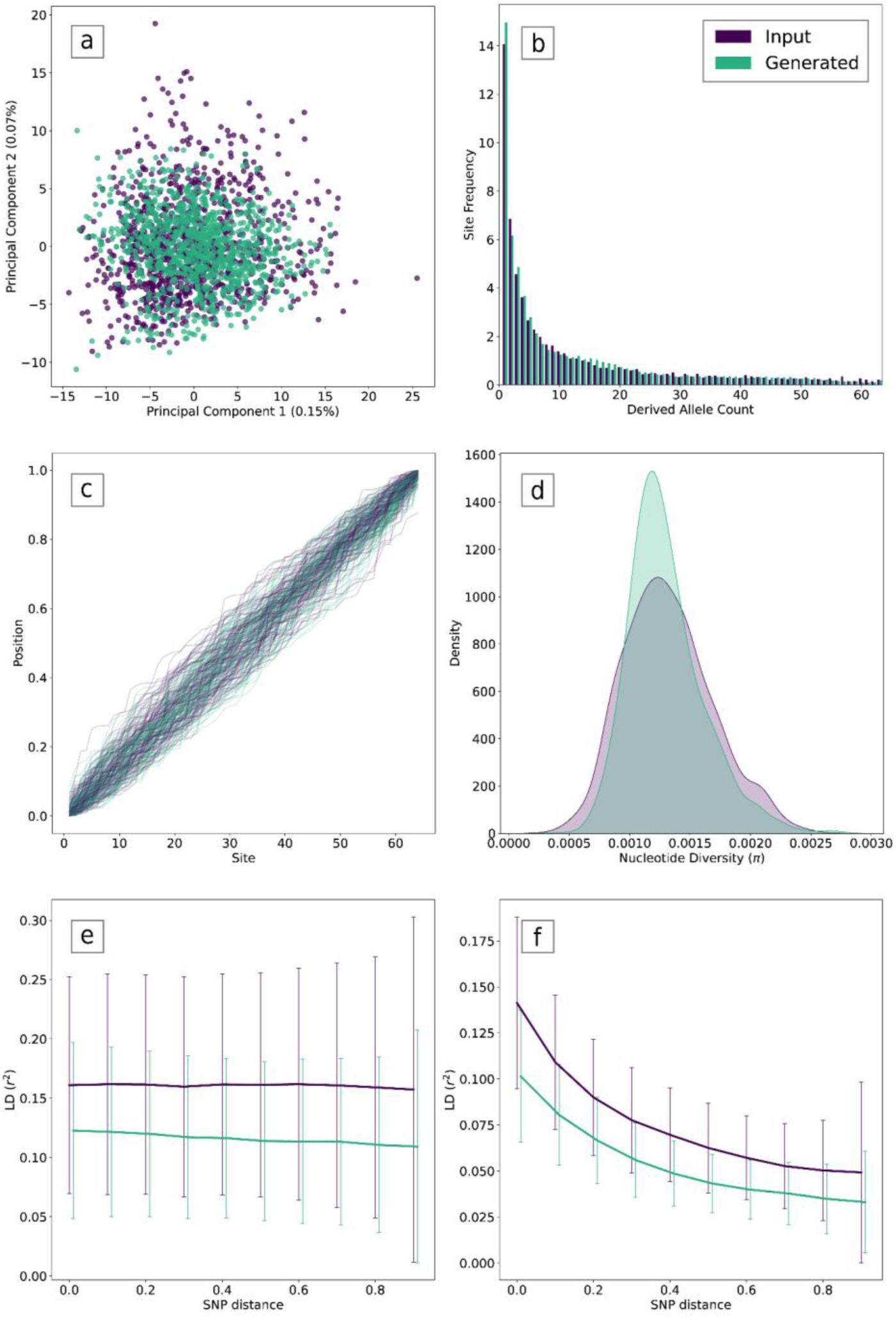
GAN performance under a standard neutral model. Input (i.e. simulated) alignments are shown in purple and generated alignments are shown in green. (a) PCA of the SFS of each alignment, calculated jointly with 100 input and generated alignments each. (b) Average SFS across 1000 simulated alignments and 1000 generated alignments. (c) Positions of polymorphisms along 100 example simulated and generated chromosomes. (d) Density plot showing the distribution of average per-site nucleotide diversity values across all input and generated alignments. (e) LD as a function of distance between polymorphisms for the non-recombinant model. (f) LD as a function of distance between polymorphisms for the recombinant model.

### Subdivided population

To test how well our GAN identifies both physical structure in the alignment and its relationship to the underlying population structure in a two-population sample, we simulated alignments under multiple two-population island models with varying levels of migration and trained our GAN on those alignments. F_ST_ between the two populations was similar between the input and generated alignments for all four migration rates examined (Fig. 3). As with LD, ^F^_ST_ is somewhat diminished in generated alignments, a pattern that may be attributable to additional noise in these alignments. We saw among the smallest 2DSWD with the two-population models as compared to other models our GAN was trained on, particularly when there was little migration (2DSWD values of 2.26 and 0.74 for migration rates of 0.1 and 1.0, respectively, with the latter being the lowest 2DSWD observed in our study; Supp. Fig. 3). This result is likely in part because these models introduce an especially clear structure into the alignments themselves— the horizontal subdivision between population one (top of the alignment) and population two (bottom of the alignment; see Fig. 4). This structure may be somewhat analogous to having distinct objects or edges in an image—something that convolutional networks were designed to detect (LeCun et al. 1998). Indeed, when we shuffled the individuals in the alignment, thereby removing that structure, our GAN was unable to recover the clear peak in the SFS at 50% derived allele frequency in the same manner as it does for the original image (Supp. Fig. 5a,c)— although it produced an abundance of alleles near 50% frequency, and was able to better mimic the density of singletons in the alignment. Interestingly, our GAN appears to recover different parts of the alignment distribution depending on whether individuals are shuffled or not (Supp. Fig. 5b,d). This may be evidence that the GAN struggles to pick up more nuanced properties of the alignments like low frequency variants when there is a very strong signal, such as a clear horizontal split in the alignment as expected in a multi-population dataset, but those properties are captured when that signal is removed. Such a result is not completely unexpected as GANs can be weak for characterizing the tails of statistical distributions (Abbasnejad et al. 2019).

**Figure 3.**
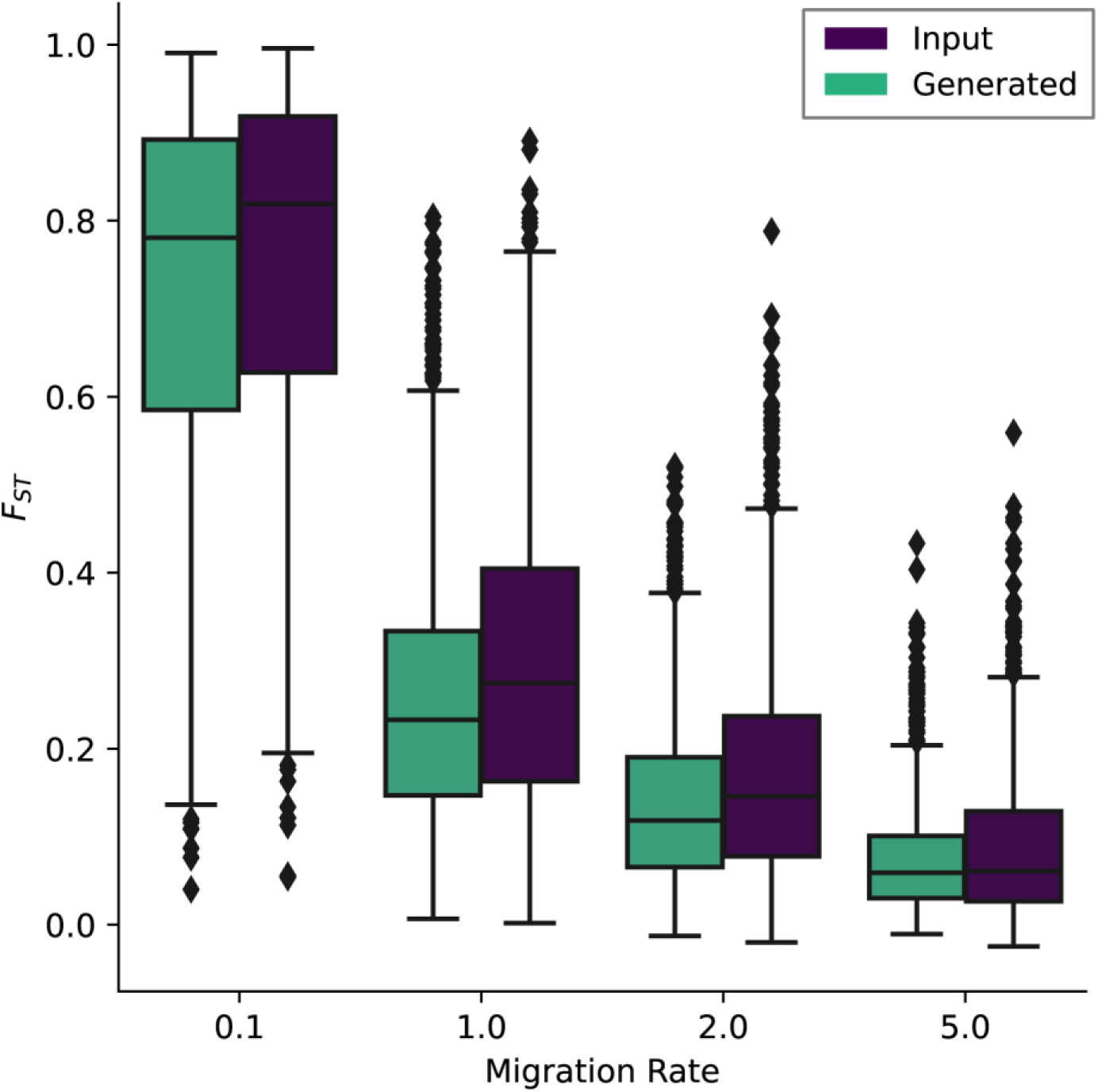
Fixation index (*F*_ST_) as a function of migration rate (shown in units of 4*Nm*) in input (purple) and generated (green) alignments. Each GAN was trained on alignments of two populations of 32 individuals and one of four bidirectional migration rates. Box and whisker plots of *F*_ST_ were generated using 1000 alignments for each migration rate.

**Figure 4.**
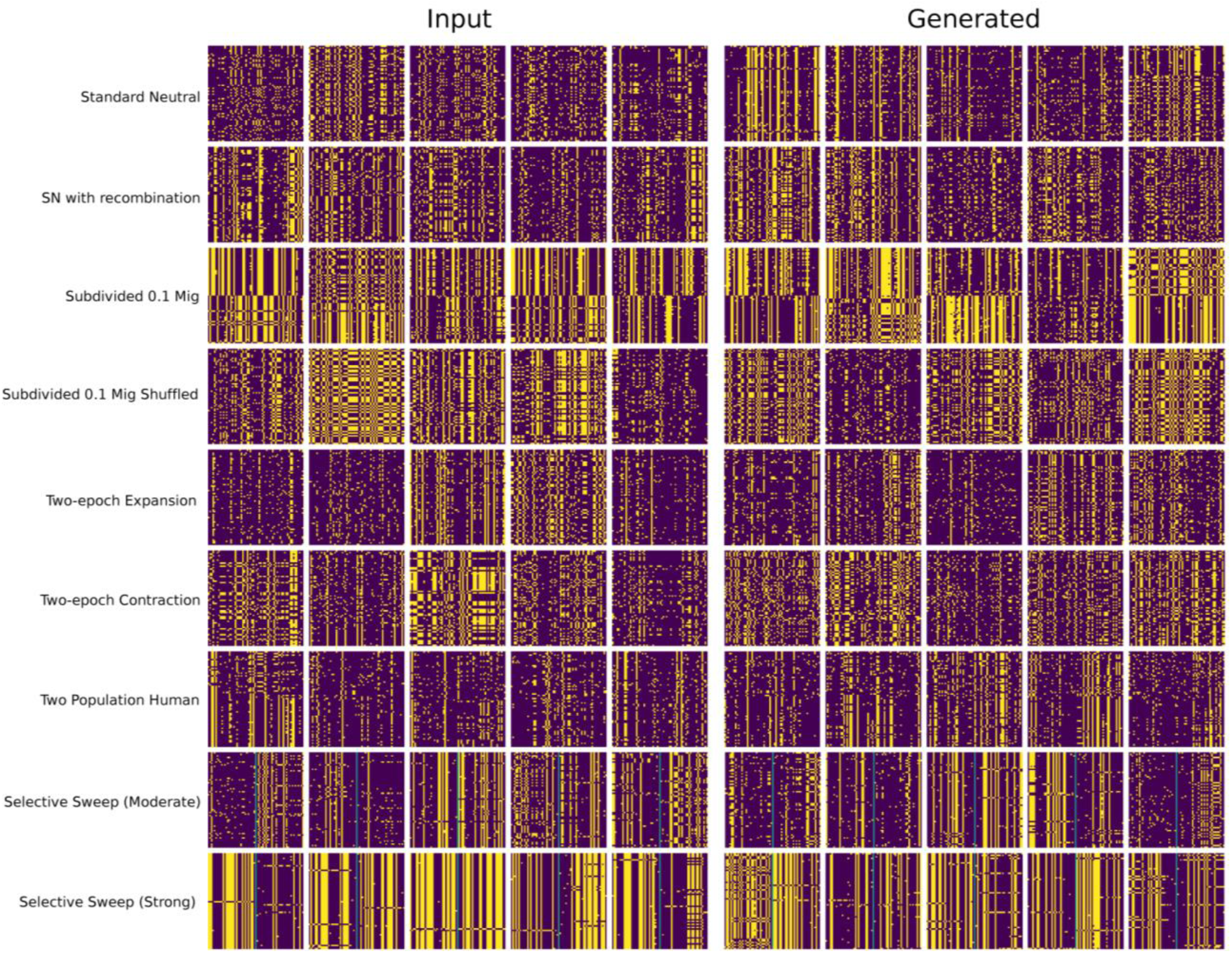
Example input (left) and generated (right) alignments for each model analyzed. Generated alignments were made using the saved network with the minimum 2DSWD. Each alignment contains 64 individuals with 64 sites. Selective sweep models (bottom two rows) show a blue line in the center of the alignment identifying the sweep site.

#### Two-epoch models of population size change

The demographic history of a population leaves a footprint in patterns of genetic variation, and accurately detecting this signature and estimating the demographic history of a population is a common analysis in population genetics (Li and Durbin 2011; Schiffels and Durbin 2014; Liu and Fu 2015, 2020; Terhorst et al. 2017). To see if our GAN can recreate this signature, we trained our GAN on alignments generated under models of population expansion and contraction and used Stairway Plot (Liu and Fu 2020) to compare the population size histories of simulated and GAN-generated alignments. Both population expansion and contraction can be mimicked by our GAN under modest population size changes (Fig. 5). Population contractions appear to be more easily captured than expansions, particularly when the magnitude of the expansion is substantial (Supp. Fig. 6). This may relate to the nature of the input data, where for histories of population expansion there is an increase in the number of rare polymorphisms (Supp. Fig. 7). Overall these alignments are more sparse and have less structure for the GAN to pick up. Additionally, any noise introduced from the GAN would have a disproportionate effect on such sparse alignments, and potentially reduce the number of rare/singleton polymorphisms.

**Figure 5.**
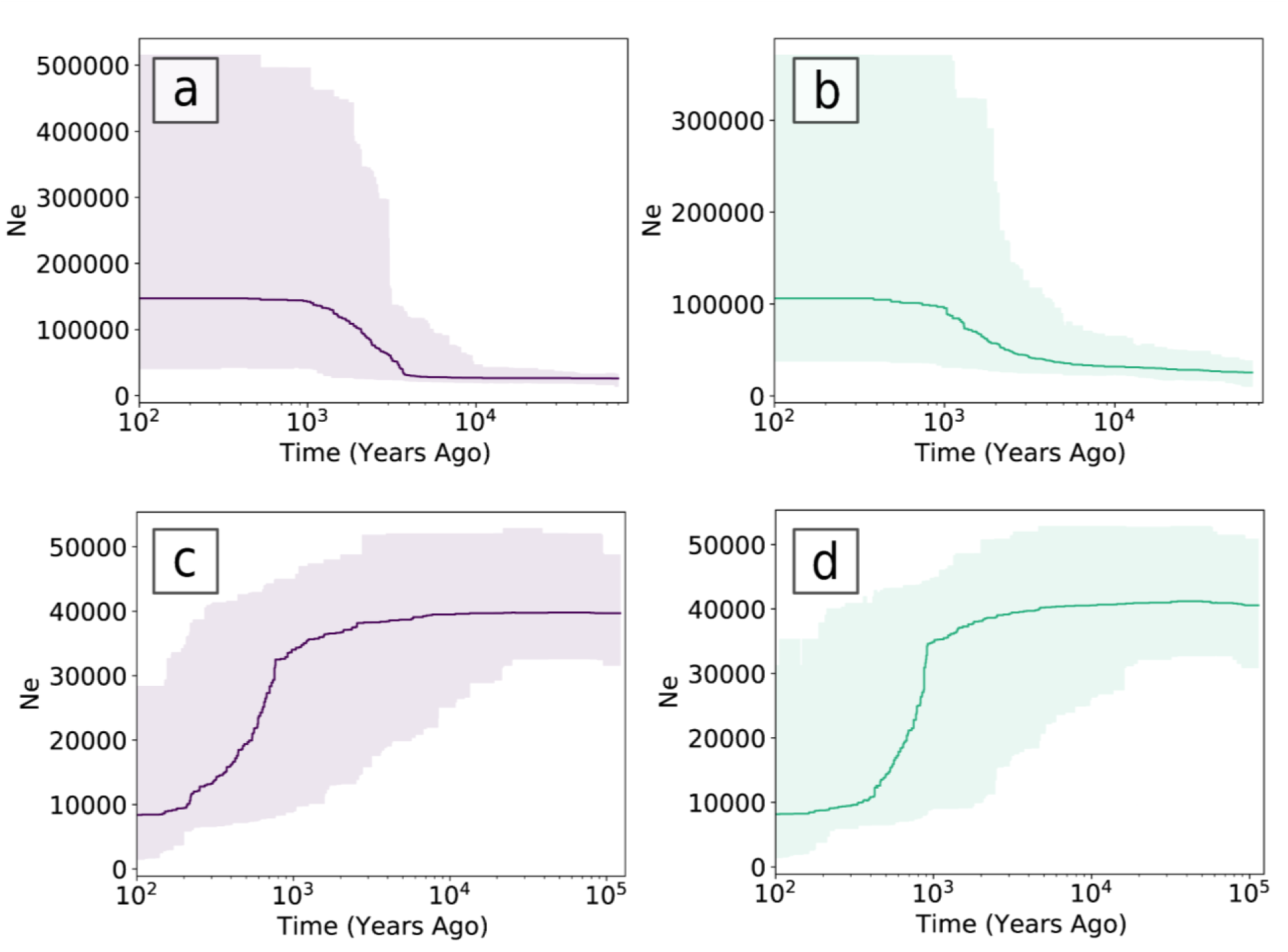
Effective population size (Ne) estimates as a function of time for input (purple) and generated (green) alignments. All estimates were calculated using Stairway Plot (version 2.1.1) with 1000 averaged site-frequency spectra and show the median and 95% CI population size estimate at each point in time. (a) Stairway Plot estimates for 10-fold population-size expansion input alignments, (b) 10-fold population-size expansion generated alignments, (c) 10-fold population contraction input alignments, (d) 10-fold population contraction generated alignments.

#### A two-population model of African and European human demographic history

Modern population genetics is increasingly more interested in identifying signatures of complex histories (Adrion et al. 2020b). To test our GAN’s ability to capture the properties of a more complex model, we trained our GAN on simulated alignments under a model of the demographic history of African and European demography (Tennessen et al. 2012). This model consists of an ancient expansion followed by a population split, two subsequent contractions of the European population, a phase of exponential expansion of the European population, and then finally rapid exponential growth in both populations continuing until the present. Interestingly, even though this is our most complex demographic model, our GAN performs perhaps better on this model than any of the other models we tested, achieving better LD estimates than neutral models (Fig. 6a) and the 2nd lowest 2DSWD of any other model (1.00 at minimum; Supp. Fig. 3). However, the 2DSWD minimum is achieved relatively early in training and the distance between generated and input alignments increases steadily after that point (Supp. Fig. 3). Also remarkably similar is the SFS of input and generated alignments for this model (Fig. 6b). Because this model includes two populations, we also investigated the joint Site-Frequency Spectrum (jSFS) produced by our GAN. Overall, our GAN is able to accurately capture the structure and relationship of the two populations in this model, with a fairly similar jSFS between the simulated and GAN-generated alignments (Fig. 6c,d). However, our GAN produces a less sharp boundary delineating alleles that are found exclusively in one population (left-most column and bottom row) or that are fixed in the European population (right-most column and top row). Although the number of sites in each of these categories is roughly similar between generated and input alignments, these strict delineations appear difficult for our GAN to capture precisely when compared to the more conspicuous delineations created by the simulations.

**Figure 6.**
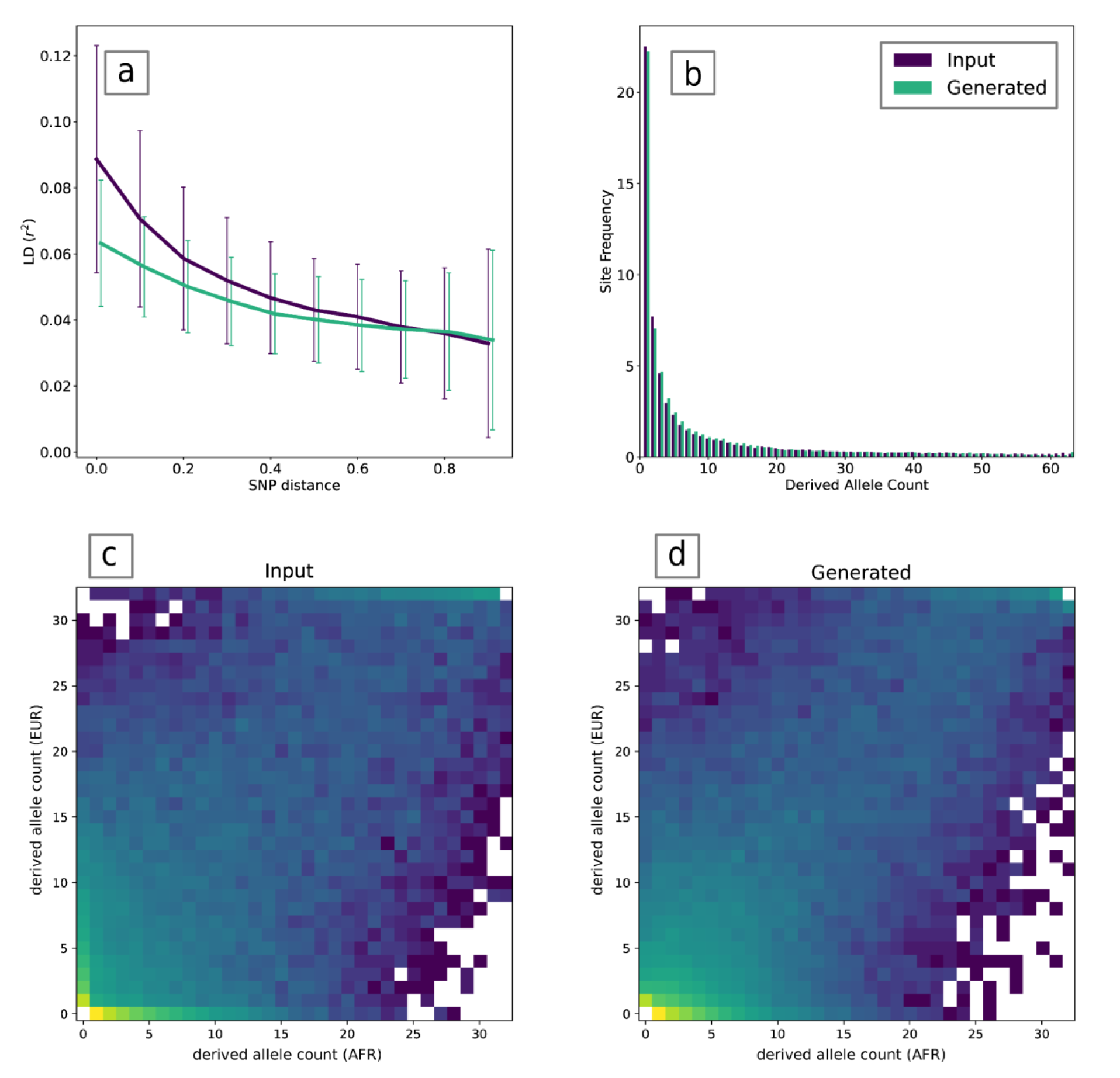
GAN results for generating alignments under the two-population out-of-Africa model from Tennessen et al. (2012). (a) LD decay as a function of distance between polymorphisms for input (purple) and generated (green) alignments. (b) Average site-frequency spectrum (SFS) over 1000 input and generated alignments. (c) joint site-frequency spectrum (jSFS) of 1000 input alignments for African (AFR) and European (EUR) individuals. (c) joint site-frequency spectrum (jSFS) of 1000 generated alignments for AFR and EUR individuals.

#### Selective sweeps

Because selective sweeps can leave a strong imprint on genetic diversity (i.e. the hitchhiking effect; (Smith and Haigh 1974; Kaplan et al. 1989; Fay and Wu 2000), but are challenging to characterize or identify (Jensen et al. 2005; Stephan 2016), we were interested in testing how well our GAN reproduces the hitchhiking effect under varying strengths of selection. Overall, our GAN was able to replicate the properties of selective sweeps but underestimated some signatures under models of strong selection. Both the moderate and strong selective sweep GANs generated alignments that show reduced nucleotide diversity around the selected site (Fig. 7) nearly identical to the pattern found in the simulated alignments. We also see a reduction in the related statistics Tajima’s D and Watterson’s θ (Supp. Fig. 8, Supp. Fig. 9). Not surprisingly, Kim and Nielsen’s ω, which seeks to detect recent hitchhiking events by taking the ratio of the amount of LD on either side of a sweep site to the amount of LD between pairs of polymorphisms on either side of the sweep site, is somewhat lower in GAN-generated than simulated alignments for strong sweeps (mean generated ω = 155.73, mean input ω = 445.29; Fig. 7). As the amount of LD produced by our GANs was lower than that produced by the corresponding simulations for each model that we examined, it is not surprising that the ω statistic deviates from the simulated distribution as well. However, it is clear that the GAN has learned the essence of the signature that ω is designed to detect: selective sweeps can produce LD blocks on either side of the sweep that do not extend across the sweep site (see example alignments, Fig. 7), and we do note that ω values produced by our GAN are similar for the moderate selection alignments (mean generated ω = 8.58, mean input ω = 8.00) and much larger than in the no-selection case (mean generated ω = 3.48, mean input ω = 3.23), especially for the GAN trained to mimic strong selection.

**Figure 7.**
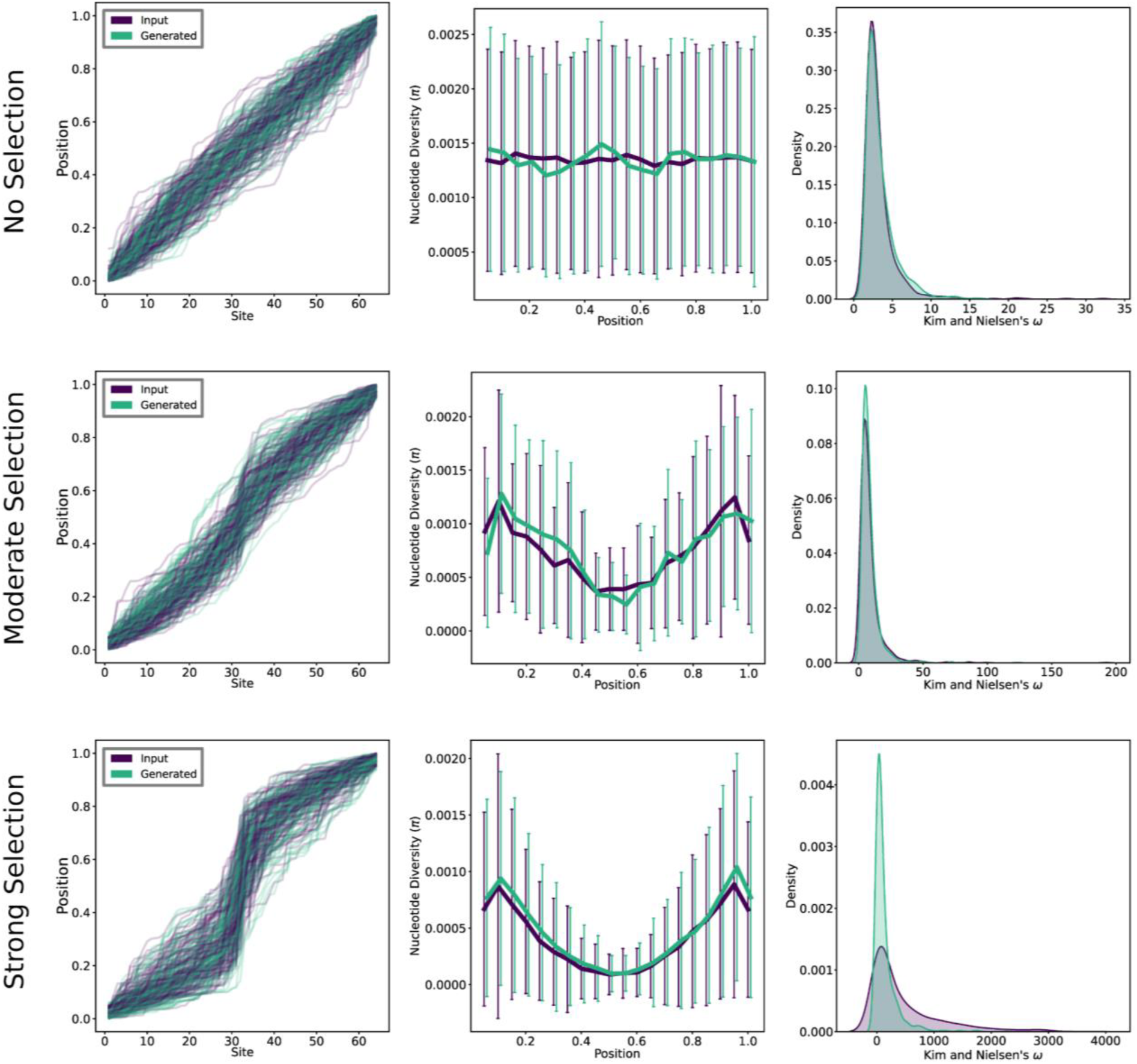
Generating selective sweeps using GANs. From top to bottom: metrics and examples from GANs trained on alignments with no selection, a moderate selective sweep, and a strong selective sweep, with the selected site occurring at the alignment midpoint. From left to right: position of each of the 64 sites in the alignment along the chromosome for input (purple) and generated (green) alignments, Average nucleotide diversity (*π*) across 1000 alignments in 0.05 position windows along the alignment (error bars represent standard deviations), and density plots showing the distribution of Kim and Nielsen’s *ω* for 1000 alignments.

### Additional Considerations and Evaluation

Throughout training and evaluation of our GAN, the potential for the GAN to generate invariant sites and the addition of noise breaking down covariances across sites provided the most consistent effects differentiating GAN-generated from simulated alignments. To quantify the effect of enforcing variation at every site in the GAN-generated alignments, we calculated several statistics to compare alignments where invariant sites were dropped from the alignment with alignments where variation was enforced (see Architecture and Implementation Methods). We compared these statistics under three models: the standard neutral model with recombination (no selection), moderate positive selection, and strong positive selection. Although simply removing invariant sites rather than enforcing variation at every site resulted in a reduction in singletons as well as poorer overlap of the generated and simulated SFS PCA (Supp. Fig. 1), enforcing variation did result in reduced LD for all models (Supp. Fig. 10). In the selective sweep models, it appears that enforcing variation results in a drop in *ω* for the model of strong selection, but looking at the moderate selection model, *ω* is actually overestimated. As such, it is unclear if the improvement to *ω* as seen in the strong selection model from dropping invariant sites is due to increased accuracy or a positive bias. Finally, there seemed to be little effect on *π* but a tendency for alignments that simply drop invariant sites to overestimate Tajima’s *D* (Supp. Fig. 10), as expected due to the reduction of singletons compared to alignments with variation enforced at every site. Ultimately, it appears that enforcing variation may have a small effect on downwardly biasing LD, but results in other improvements that make GAN generated alignments more similar to the input alignments.

To quantify the impact of noise introduced by the GAN on its ability to recreate the properties of the input alignments, we compared input and GAN-generated alignments to input alignments where a random one percent of values were flipped. We performed this experiment for the same models examined above when testing the impact of enforcing variation. As predicted, the added noise resulted in reduced LD across all models and lower *ω* for models of selection (Supp. Fig. 11). Added noise had little effect on *π*, but did appear to positively influence Tajima’s *D* across most windows for all models. Interestingly, this small percentage of added noise had a larger effect on LD, *ω*, and Tajima’s D for selection models, indicating that the GAN generated alignments come remarkably close to capturing the statistics of the input alignments, but for models with reduced diversity and stronger correlations across sites (such as in selection models), even small amounts of noise can have noticeable effects. We also investigated the effect of noise on *^F^*_ST_ for subdivided population models (Supp. Fig. 12). As in the selection models, added noise reliably reduced *F*_ST_ between populations to levels similar to those observed in GAN-generated alignments.

## DISCUSSION

In the present work we have demonstrated the efficacy of using GANs to generate artificial population genetic alignments. After testing several architectures, we were able to design a GAN that could accurately capture and replicate the properties of alignments under a variety of evolutionary models. For many of these properties, such as nucleotide diversity and the position of sites along the alignment, generated alignments from our GAN were largely indistinguishable from the input alignments. For others, such as measures of LD, even though the GAN-generated alignments produced values differing from the simulated distribution, the qualitative dynamics of these statistics reflected those of the simulated population genetic model (e.g. decaying LD as a function of the distance between polymorphisms for recombinant models, and the presence of distinct LD blocks on either side of a selective sweep).

Although our GAN performed well for most models tested, it underperformed in some consistent ways. Most notably was the addition of noise or less distinct boundaries in generated alignments as compared to the simulated alignments. This effect predictably resulted in the reduction of certain metrics that rely on structure between individuals or sites (e.g. LD and *^F^*_ST_), something also seen in previous genetic GANs (Wang *et al*. 2020; Yelmen *et al*. 2021). As well, although the number of variable sites was close between simulated and generated alignments, we were unable to ensure variation at every site using just the GAN architecture, and instead enforced variation by sampling an individual in proportion to its allele probabilities at otherwise monomorphic sites. While inaccuracy due to noise is most proximally due to inadequate GAN performance, the ultimate cause of all of these discrepancies results from comparing an exact and specific population genetic alignment from simulations to an alignment that is not exact by nature. Simulators produce alignments that guarantee the presence of variation at every segregating site, and the models that produce linkage disequilibrium and population structure do not introduce ‘noise’ beyond that governed by the user-specified model parameters themselves (e.g. a migration rate parameter that erodes population structure). Although improvements to the architecture and the use of more sophisticated GANs may produce generated alignments that more accurately capture these properties, unless the exact function to produce an alignment for a given model is identified by the network GAN generated alignments will always be an inexact approximation as compared to simulated alignments.

As presented here, our GAN provides a proof-of-concept for a population genetic GAN in the nature of GANs as broadly implemented in the deep learning literature: unlike previously developed population genetic GANs, ours is fully-differentiable (in terms of the density distribution of alignments) and generates sequences for the whole population sample at once. This design implies that, while our GAN can be useful for more traditional uses, such as data augmentation, it perhaps more importantly provides a basis with which to explore more advanced applications of GANs to population genetics.

One potential application with which to extend this work that may prove to be particularly useful to population genetics is style transfer (Karras *et al*. 2018), where subproperties of the input are identified by the network and can then be transferred to other data where that subproperty is absent. In a population genetic context, the ability to transfer styles from one source to another could be used to ‘transfer’ one evolutionary process, such as positive selection, to a population genetic alignment where selection is not believed to have occurred. In this example, the positive selection ‘style’ can be learned from simulated data, while the examples without sweeps can use empirical data that includes all of the challenges such data may present for detecting selection (e.g. unknown true demographic history, cryptic population structure, low data quality, etc). Thus, this approach presents the possibility of combining the strengths of reductionist generative models with those of GANs, and could potentially be used to produce more accurate training data for machine learning-based methods for detecting selective sweeps (Pybus *et al*. 2015; Schrider and Kern 2016; Kern and Schrider 2018; Whitehouse and Schrider 2022). Such uses may prove to be one of the strengths of GANs for population genetic data over using simulators alone, as they provide ability to capture and recreate subtle properties of the data that cannot be encompassed by simple idealized models used for simulations. Similarly, one could take alignments from empirical data and transfer various demographic hypotheses to those alignments to help generate the reference table for performing approximate Bayesian computation. Much like transferring selection ‘styles’, this example provides the obvious benefit of not requiring pre-defined population parameters for generating all aspects of the desired training data. Finally, a particularly intriguing use would involve combining work on creating an interpretable latent space, where population parameters can be translated into different parts of a multi-dimensional Gaussian (Liu *et al*. 2022), with the ability to project generated examples onto this latent space (Karras *et al*. 2021). While the latent space interpretation would allow for a simulator that is instantaneous once trained, the combination with latent space projection would make the network capable of inferring the demographic parameters of real data and testing evolutionary hypotheses using the network directly.

Ultimately, this work and its potential expansions further demonstrate the potential of deep-learning methods for population genetic research. While some may deride deep-learning as inferior to more traditional statistical methods due to their (current) black box nature, we urge skeptical readers to momentarily suspend this view. Instead, we implore readers to explore the continually-expanding collection of deep-learning networks alongside their applications, and consider how these tools could be used to help facilitate advances in their own systems and fields. As we have shown here, researchers do not need to start from scratch and can repurpose existing networks for evolutionary applications. Unsurprisingly, such repurposing has led to several creative purposes well outside of the developer’s initial intention, such as applying cycleGANs (Zhu *et al*. 2017) to normalize single cell RNA-seq data across datasets (Khan *et al*. 2022) and using natural language processing models to predict the effect of nucleotide variants (Benegas *et al*. 2022). At a time when unprecedented resources are being used to develop deep-learning technologies outside of biology, biologists from all fields should consider the myriad ways in which these technologies can be leveraged to solve problems and answer longstanding questions.

## DATA AVAILABILITY

Scripts and additional example images available at github.com/SchriderLab/PG-Alignments-GAN

## ACKNOWLEDGEMENTS

We would like to thank Andrew Kern for comments on the manuscript, Nick Matthew for his work and guidance on GANs prior to the development of this project, as well as other members of our lab: Logan Whitehouse, Amjad Dabi, Becca Love, and Anton Suvrov for their advice and assistance.

