## Supplemental File 1 for "This population does not exist: learning the distribution of evolutionary histories with generative adversarial networks"

### Supporting Information

**Supplemental Table 1.** Models used for this study and their commands for simulation.

| | |
|---|---|
| Standard neutral | ms 64 20000 -s 64 |
| Standard neutral with recombination | ms 64 20000 -s 64 -r 13.6 1000 |
| Subdivided | ms 64 20000 -s 64 -I 2 32 32 0.1<br>ms 64 20000 -s 64 -I 2 32 32 1.0<br>ms 64 20000 -s 64 -I 2 32 32 2.0<br>ms 64 20000 -s 64 -I 2 32 32 5.0 |
| Two-epoch contraction | ms 64 20000 -t 40 -eN 0 0.5 -eN 0.01 5 |
| Two-epoch expansion | ms 64 20000 -t 100 -eN 0 5 -eN 0.01 0.5 |
| Two-population Out-of-Africa | species = stdpopsim.get_species("HomSap")<br>model = species.get_demographic_model("OutOfAfrica_2T12")<br>contig = species.get_contig("chr22", length_multiplier=0.001)<br>samples = model.get_samples(32,32)<br>engine = stdpopsim.get_engine("msprime")<br>ts = engine.simulate(model, contig, samples) |
| Selective sweep (strong) | discoal 64 20000 1000 -t 125 -ws 0 -x 0.5 -r 125 -a 2000 |
| Selective sweep (moderate) | discoal 64 20000 1000 -t 45 -r 45 -ws 0 -x 0.5 -a 100 |

**Supplemental Table 2.** Sorted results from hyper-parameter tuning. The 2D sliced Wasserstein distance (2DSWD) between the distribution of input and generated SFS PCAs was calculated after every 100 epochs for 10,000 epochs and averaged across the last 5 100 epoch iterations. Hypertuning results are for various hyperparameter values used to train a GAN on 10,000 alignments generated under a standard neutral simulation. The final chosen set of hyperparameters is shown at the top of the table, with the smallest 2DSWD.

| **2DSWD** | **Critic Iterations** | **Negative Slope** | **Batch Size** | **Learning Rate** | **Lambda** |
|---|---|---|---|---|---|
| 5.87411 | 5 | 0.1 | 256 | 0.0002 | 10 |
| 5.93474 | 5 | 0.05 | 128 | 0.00005 | 7 |
| 5.94295 | 3 | 0.2 | 64 | 0.0001 | 7 |
| 6.00638 | 3 | 0.2 | 128 | 0.0002 | 7 |
| 6.04408 | 3 | 0.1 | 256 | 0.00005 | 10 |
| 6.05793 | 3 | 0.2 | 256 | 0.00005 | 7 |
| 6.10092 | 3 | 0.1 | 128 | 0.0002 | 7 |
| 6.10765 | 3 | 0.1 | 128 | 0.0001 | 7 |
| 6.17229 | 5 | 0.05 | 256 | 0.0001 | 10 |
| 6.1919 | 5 | 0.05 | 128 | 0.0002 | 10 |
| 6.19733 | 5 | 0.1 | 64 | 0.0002 | 10 |
| 6.19958 | 5 | 0.1 | 64 | 0.00005 | 10 |
| 6.20759 | 5 | 0.1 | 256 | 0.0001 | 10 |
| 6.229 | 3 | 0.1 | 64 | 0.00005 | 13 |

| | | | | | |
|---|---|---|---|---|---|
| 6.26189 | 3 | 0.05 | 64 | 0.0001 | 13 |
| 6.262 | 3 | 0.1 | 128 | 0.0001 | 13 |
| 6.27331 | 3 | 0.2 | 128 | 0.0001 | 7 |
| 6.27816 | 3 | 0.2 | 256 | 0.0001 | 10 |
| 6.2905 | 3 | 0.1 | 256 | 0.0002 | 10 |
| 6.29516 | 3 | 0.05 | 64 | 0.00005 | 7 |
| 6.30886 | 3 | 0.2 | 64 | 0.00005 | 7 |
| 6.32172 | 3 | 0.05 | 256 | 0.0002 | 7 |
| 6.33799 | 5 | 0.05 | 64 | 0.0002 | 13 |
| 6.34701 | 3 | 0.2 | 256 | 0.0002 | 13 |
| 6.36392 | 5 | 0.05 | 64 | 0.0002 | 10 |
| 6.36868 | 5 | 0.1 | 128 | 0.0001 | 7 |
| 6.37928 | 3 | 0.2 | 256 | 0.0001 | 7 |
| 6.38694 | 3 | 0.1 | 64 | 0.0002 | 13 |
| 6.40434 | 5 | 0.05 | 128 | 0.0002 | 7 |
| 6.40612 | 5 | 0.2 | 64 | 0.0001 | 7 |
| 6.41192 | 5 | 0.05 | 64 | 0.00005 | 13 |

| | | | | | |
|---|---|---|---|---|---|
| 6.4121 | 5 | 0.2 | 128 | 0.0001 | 10 |
| 6.41471 | 3 | 0.1 | 256 | 0.0002 | 7 |
| 6.42442 | 3 | 0.1 | 256 | 0.0002 | 13 |
| 6.42818 | 3 | 0.05 | 128 | 0.0002 | 13 |
| 6.45462 | 3 | 0.1 | 64 | 0.0002 | 7 |
| 6.45554 | 3 | 0.2 | 64 | 0.0002 | 13 |
| 6.45769 | 3 | 0.1 | 128 | 0.00005 | 13 |
| 6.46328 | 5 | 0.1 | 64 | 0.0001 | 7 |
| 6.46528 | 5 | 0.1 | 64 | 0.0001 | 10 |
| 6.48234 | 5 | 0.1 | 128 | 0.0002 | 7 |
| 6.48584 | 5 | 0.2 | 128 | 0.0002 | 7 |
| 6.48753 | 5 | 0.2 | 64 | 0.00005 | 10 |
| 6.49694 | 5 | 0.1 | 256 | 0.0002 | 13 |
| 6.50457 | 5 | 0.05 | 256 | 0.00005 | 10 |
| 6.50636 | 5 | 0.2 | 256 | 0.00005 | 7 |
| 6.50718 | 5 | 0.1 | 256 | 0.0001 | 7 |
| 6.51397 | 3 | 0.1 | 128 | 0.00005 | 10 |

| | | | | | |
|---|---|---|---|---|---|
| 6.52245 | 5 | 0.05 | 128 | 0.00005 | 13 |
| 6.53101 | 5 | 0.05 | 128 | 0.00005 | 10 |
| 6.55597 | 3 | 0.2 | 128 | 0.0001 | 10 |
| 6.56345 | 3 | 0.2 | 128 | 0.0002 | 13 |
| 6.58064 | 5 | 0.05 | 64 | 0.00005 | 7 |
| 6.58214 | 5 | 0.1 | 128 | 0.0001 | 13 |
| 6.59873 | 5 | 0.05 | 64 | 0.0001 | 10 |
| 6.59877 | 5 | 0.1 | 128 | 0.0002 | 10 |
| 6.60062 | 3 | 0.05 | 64 | 0.0001 | 7 |
| 6.6028 | 3 | 0.2 | 256 | 0.0002 | 7 |
| 6.62975 | 5 | 0.05 | 64 | 0.0002 | 7 |
| 6.64012 | 3 | 0.2 | 128 | 0.0001 | 13 |
| 6.64276 | 5 | 0.2 | 64 | 0.0001 | 13 |
| 6.65617 | 3 | 0.1 | 128 | 0.0002 | 10 |
| 6.66339 | 5 | 0.05 | 256 | 0.00005 | 13 |
| 6.66821 | 5 | 0.1 | 128 | 0.00005 | 10 |
| 6.67048 | 3 | 0.05 | 128 | 0.0002 | 7 |

Population Genetic Alignment GAN

| | | | | | |
|---|---|---|---|---|---|
| 6.69036 | 5 | 0.05 | 256 | 0.0001 | 7 |
| 6.69057 | 5 | 0.05 | 128 | 0.0001 | 13 |
| 6.698 | 3 | 0.2 | 64 | 0.0001 | 13 |
| 6.6985 | 3 | 0.2 | 64 | 0.00005 | 13 |
| 6.70101 | 5 | 0.2 | 64 | 0.0002 | 10 |
| 6.70287 | 3 | 0.1 | 128 | 0.00005 | 7 |
| 6.70348 | 5 | 0.05 | 128 | 0.0002 | 13 |
| 6.70428 | 5 | 0.2 | 128 | 0.00005 | 13 |
| 6.7089 | 3 | 0.1 | 256 | 0.0001 | 7 |
| 6.72157 | 5 | 0.05 | 256 | 0.0002 | 7 |
| 6.72375 | 3 | 0.05 | 256 | 0.00005 | 7 |
| 6.72785 | 5 | 0.2 | 64 | 0.0002 | 7 |
| 6.74168 | 3 | 0.05 | 256 | 0.0001 | 10 |
| 6.7419 | 5 | 0.05 | 256 | 0.0002 | 10 |
| 6.74266 | 3 | 0.1 | 64 | 0.00005 | 10 |
| 6.74598 | 5 | 0.2 | 64 | 0.00005 | 13 |
| 6.74975 | 3 | 0.05 | 128 | 0.00005 | 10 |

| | | | | | |
|---|---|---|---|---|---|
| 6.76969 | 5 | 0.05 | 64 | 0.00005 | 10 |
| 6.77618 | 5 | 0.1 | 64 | 0.00005 | 7 |
| 6.77886 | 3 | 0.05 | 256 | 0.0001 | 13 |
| 6.78035 | 5 | 0.1 | 128 | 0.0002 | 13 |
| 6.79024 | 3 | 0.05 | 128 | 0.00005 | 7 |
| 6.79112 | 3 | 0.1 | 64 | 0.0001 | 10 |
| 6.79691 | 3 | 0.1 | 64 | 0.0001 | 13 |
| 6.79903 | 5 | 0.2 | 256 | 0.00005 | 10 |
| 6.80702 | 3 | 0.1 | 64 | 0.0002 | 10 |
| 6.80986 | 3 | 0.2 | 64 | 0.00005 | 10 |
| 6.81372 | 3 | 0.1 | 64 | 0.0001 | 7 |
| 6.83414 | 3 | 0.2 | 64 | 0.0002 | 7 |
| 6.83985 | 5 | 0.2 | 256 | 0.0001 | 13 |
| 6.84113 | 3 | 0.2 | 256 | 0.0001 | 13 |
| 6.84245 | 5 | 0.1 | 64 | 0.0002 | 7 |
| 6.85644 | 3 | 0.1 | 256 | 0.00005 | 7 |
| 6.87124 | 3 | 0.2 | 64 | 0.0002 | 10 |

| | | | | | |
|---|---|---|---|---|---|
| 6.87158 | 5 | 0.2 | 256 | 0.0001 | 7 |
| 6.87834 | 5 | 0.2 | 64 | 0.0001 | 10 |
| 6.88091 | 5 | 0.2 | 128 | 0.00005 | 10 |
| 6.88179 | 5 | 0.2 | 64 | 0.00005 | 7 |
| 6.89394 | 5 | 0.1 | 256 | 0.0001 | 13 |
| 6.91445 | 3 | 0.05 | 128 | 0.0001 | 13 |
| 6.92013 | 3 | 0.05 | 64 | 0.0002 | 10 |
| 6.92505 | 5 | 0.2 | 256 | 0.0001 | 10 |
| 6.92947 | 3 | 0.05 | 256 | 0.0002 | 10 |
| 6.94061 | 3 | 0.1 | 64 | 0.00005 | 7 |
| 6.94177 | 3 | 0.05 | 128 | 0.00005 | 13 |
| 6.94303 | 5 | 0.1 | 256 | 0.0002 | 7 |
| 6.94823 | 5 | 0.1 | 64 | 0.0001 | 13 |
| 6.96352 | 3 | 0.1 | 128 | 0.0002 | 13 |
| 6.97825 | 5 | 0.2 | 64 | 0.0002 | 13 |
| 6.98608 | 5 | 0.1 | 128 | 0.0001 | 10 |
| 6.99399 | 3 | 0.05 | 64 | 0.0002 | 13 |

| | | | | | |
|---|---|---|---|---|---|
| 6.99967 | 3 | 0.05 | 256 | 0.00005 | 10 |
| 7.00575 | 5 | 0.1 | 64 | 0.0002 | 13 |
| 7.01481 | 5 | 0.05 | 256 | 0.0001 | 13 |
| 7.02492 | 3 | 0.2 | 256 | 0.00005 | 10 |
| 7.03051 | 5 | 0.2 | 128 | 0.0001 | 13 |
| 7.03168 | 3 | 0.2 | 256 | 0.00005 | 13 |
| 7.04809 | 5 | 0.2 | 256 | 0.0002 | 13 |
| 7.05967 | 5 | 0.05 | 64 | 0.0001 | 7 |
| 7.06284 | 3 | 0.05 | 256 | 0.0001 | 7 |
| 7.06631 | 3 | 0.2 | 256 | 0.0002 | 10 |
| 7.0688 | 3 | 0.05 | 128 | 0.0002 | 10 |
| 7.09667 | 3 | 0.05 | 64 | 0.0002 | 7 |
| 7.1083 | 5 | 0.05 | 64 | 0.0001 | 13 |
| 7.12276 | 3 | 0.05 | 128 | 0.0001 | 7 |
| 7.12441 | 5 | 0.1 | 256 | 0.00005 | 10 |
| 7.15015 | 3 | 0.2 | 128 | 0.0002 | 10 |
| 7.15558 | 3 | 0.1 | 256 | 0.0001 | 13 |

| | | | | | |
|---|---|---|---|---|---|
| 7.15796 | 5 | 0.2 | 128 | 0.0002 | 10 |
| 7.16688 | 5 | 0.05 | 128 | 0.0001 | 10 |
| 7.16808 | 3 | 0.2 | 64 | 0.0001 | 10 |
| 7.18039 | 3 | 0.2 | 128 | 0.00005 | 10 |
| 7.19611 | 5 | 0.2 | 256 | 0.00005 | 13 |
| 7.1972 | 3 | 0.05 | 64 | 0.00005 | 13 |
| 7.2513 | 3 | 0.05 | 256 | 0.00005 | 13 |
| 7.25356 | 3 | 0.05 | 64 | 0.0001 | 10 |
| 7.26264 | 3 | 0.1 | 128 | 0.0001 | 10 |
| 7.27577 | 5 | 0.2 | 128 | 0.00005 | 7 |
| 7.30135 | 5 | 0.1 | 256 | 0.00005 | 13 |
| 7.32328 | 5 | 0.05 | 256 | 0.00005 | 7 |
| 7.33035 | 5 | 0.1 | 64 | 0.00005 | 13 |
| 7.34931 | 3 | 0.1 | 256 | 0.0001 | 10 |
| 7.36709 | 3 | 0.2 | 128 | 0.00005 | 7 |
| 7.3816 | 5 | 0.1 | 256 | 0.00005 | 7 |
| 7.39383 | 5 | 0.05 | 128 | 0.0001 | 7 |

| | | | | | |
|---|---|---|---|---|---|
| 7.40595 | 5 | 0.2 | 256 | 0.0002 | 10 |
| 7.44567 | 5 | 0.2 | 256 | 0.0002 | 7 |
| 7.44807 | 5 | 0.2 | 128 | 0.0002 | 13 |
| 7.48629 | 5 | 0.1 | 128 | 0.00005 | 7 |
| 7.53794 | 3 | 0.2 | 128 | 0.00005 | 13 |
| 7.5714 | 5 | 0.2 | 128 | 0.0001 | 7 |
| 7.62356 | 3 | 0.05 | 128 | 0.0001 | 10 |
| 7.63378 | 3 | 0.05 | 256 | 0.0002 | 13 |
| 7.80113 | 3 | 0.1 | 256 | 0.00005 | 13 |
| 7.8211 | 5 | 0.05 | 256 | 0.0002 | 13 |
| 7.85496 | 5 | 0.1 | 128 | 0.00005 | 13 |
| 8.71081 | 3 | 0.05 | 64 | 0.00005 | 10 |

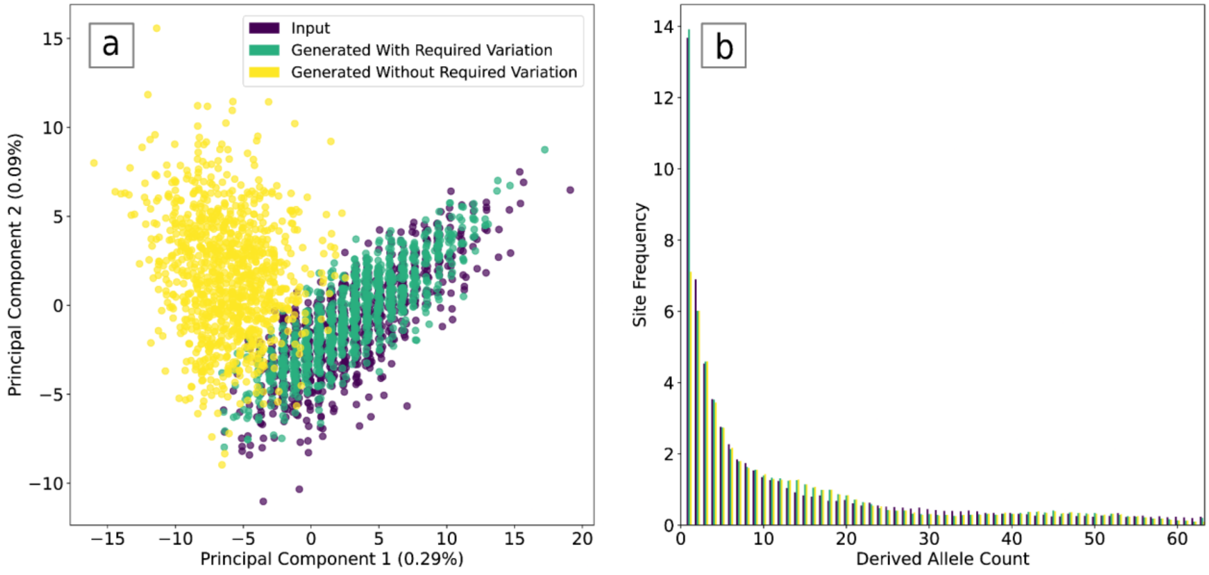


**Supplemental Figure 1.** GAN performance when invariant sites are dropped from the alignment compared (without required variation) to alignments where variation is forced through weighted random sampling (with required variation). (a) PCA of the site-frequency spectrum comparing 200 individual input alignments to GAN generated alignments with and without required variation. (b) Average site-frequency spectrum across 1000 alignments for input alignments and GAN generated alignments with and without required variation.

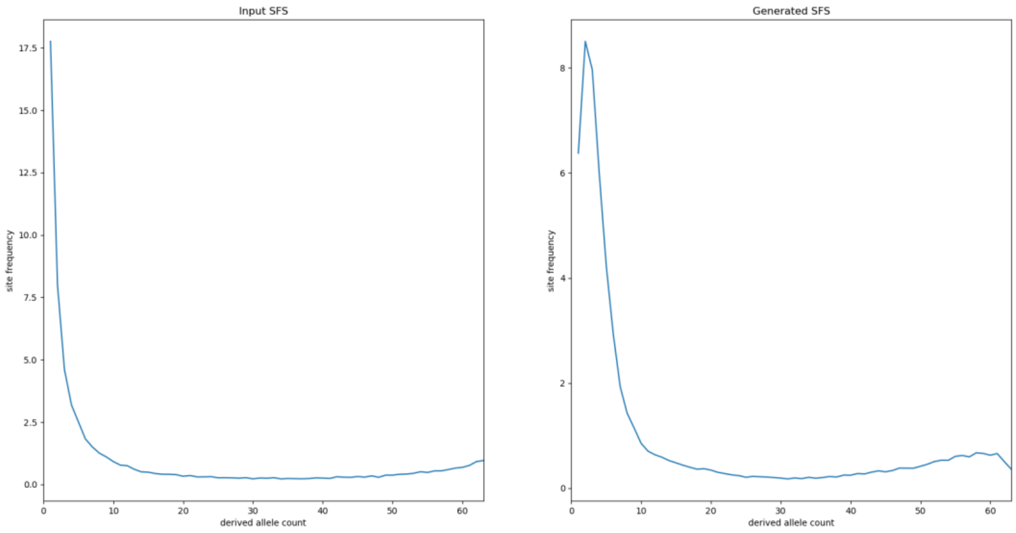


**Supplemental Figure 2.** Average site-frequency spectrum from 1000 input (left) and generated (right) alignments using a custom loss penalty on monomorphic sites (c~~C~~ustom penalty shown in Eq. 2).

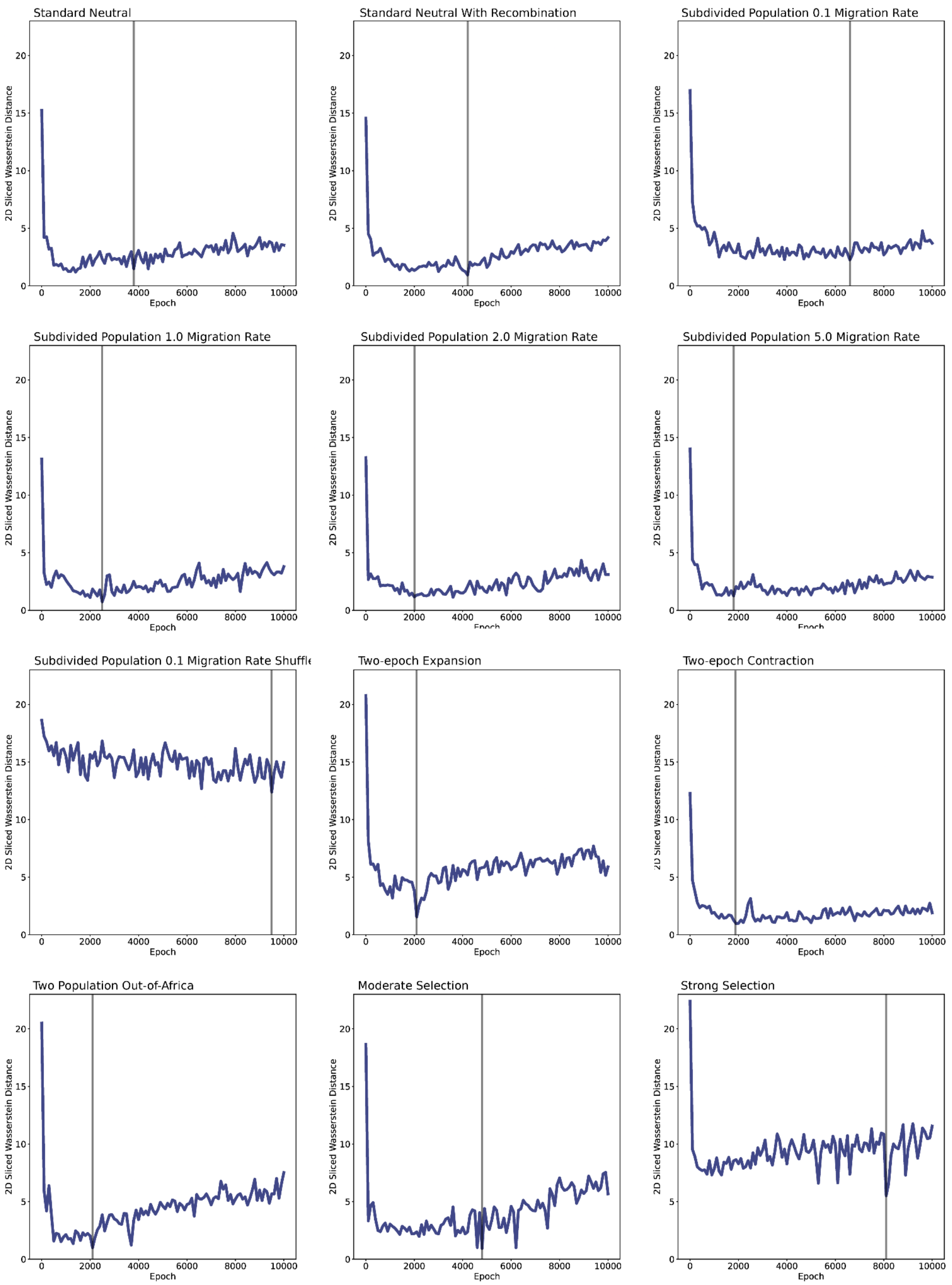


**Supplemental Figure 3.** 2D sliced Wasserstein distance (2DSWD) between the PCA values calculated from input and generated site-frequency spectra throughout training. Vertical line represents the 2DSWD minimum value during training used for evaluation.

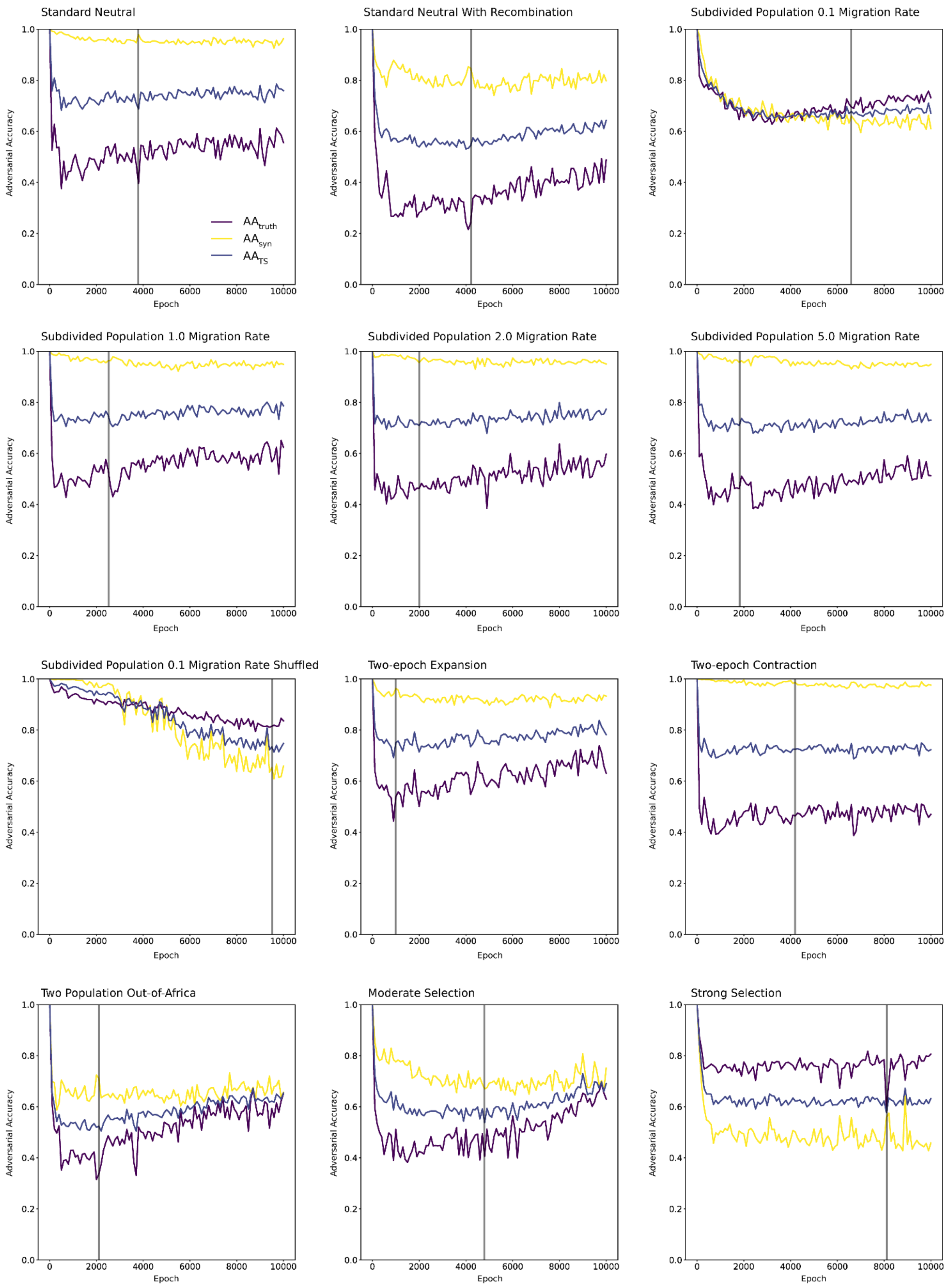


**Supplemental Figure 4.** Nearest neighbor adversarial accuracy (AA) scores throughout training. Vertical line represents the 2DSWD minimum value during training used for evaluation.

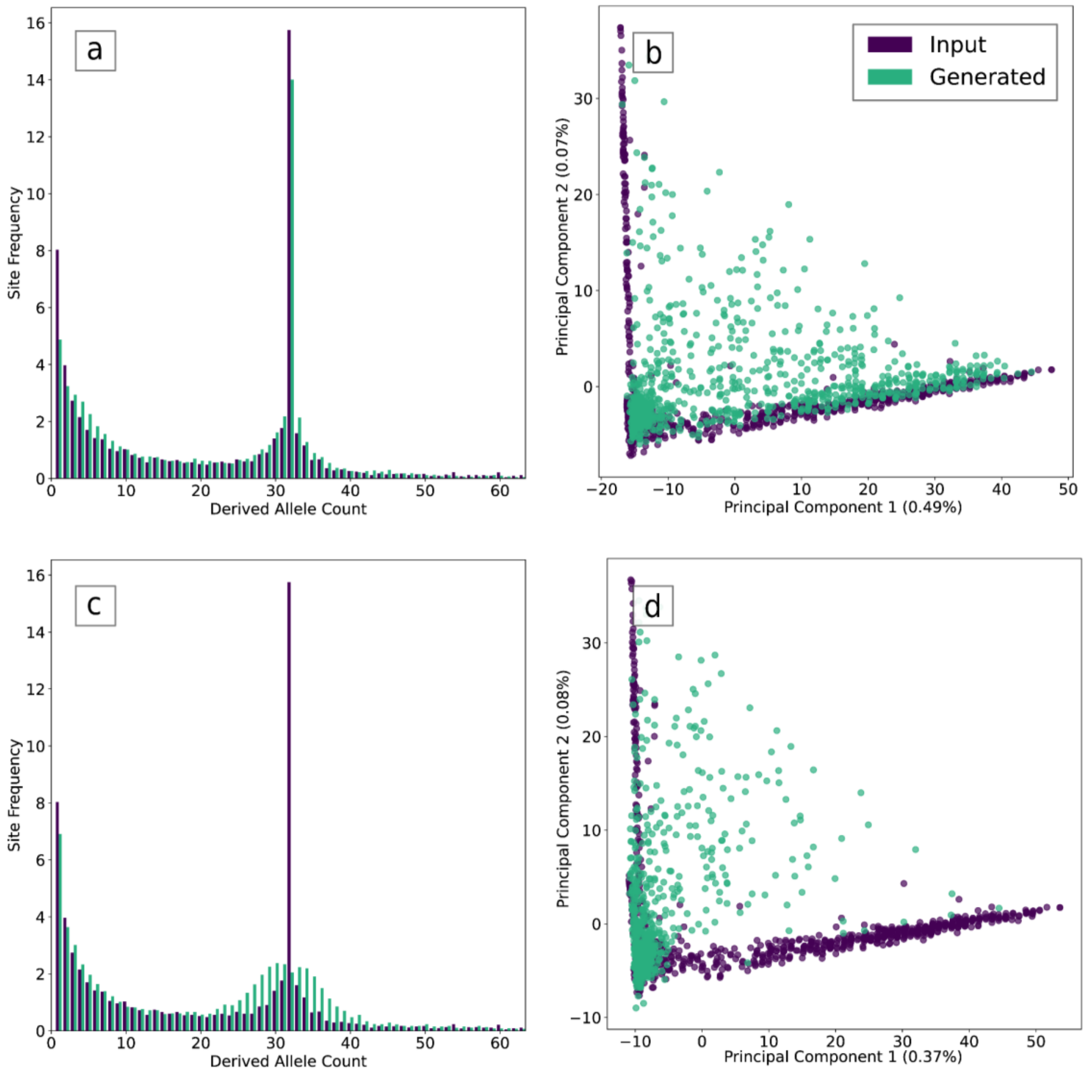


**Supplemental Figure 5.** GAN performance on unshuffled (a,b) and shuffled (c,d) structured population alignments with a 0.1 migration rate. (a) Average SFS of 1,000 input (purple) and GAN generated (green) alignments of an unshuffled structured population. (b) PCA of the SFS of 200 input (purple) and GAN generated (green) alignments of an unshuffled structured population. (c) Average SFS of 1,000 input (purple) and GAN generated (green) alignments of a shuffled structured population. (d) PCA of the SFS of 200 input (purple) and GAN generated (green) alignments of a shuffled structured population.

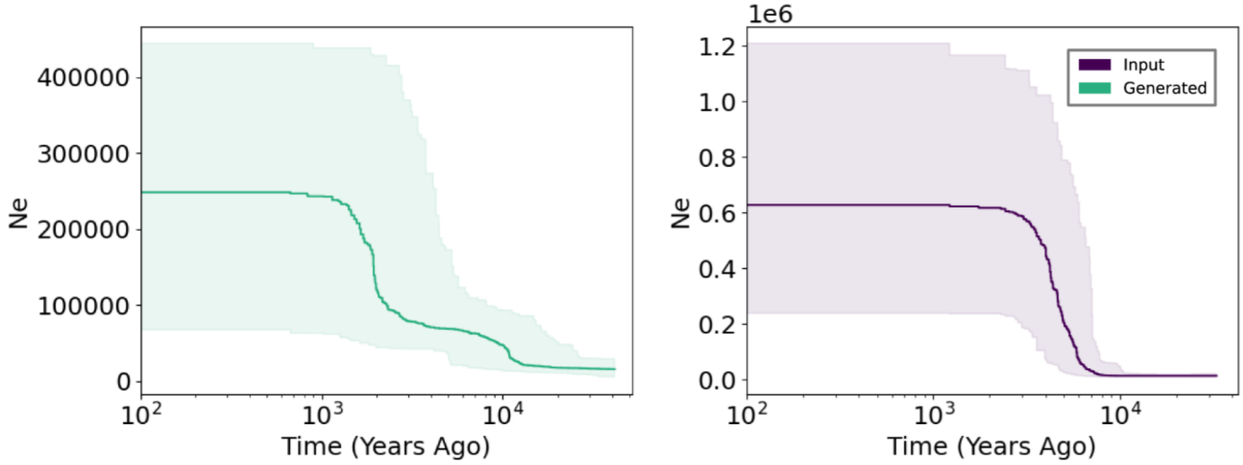


**Supplemental Figure 6.** Effective population size (Ne) estimates for a population experiencing a 50-fold increase as a function of time for input (purple) and generated (green) alignments. All estimates were calculated using Stairwayplot (version 2.1.1) and show the median and 95% CI population size estimate.

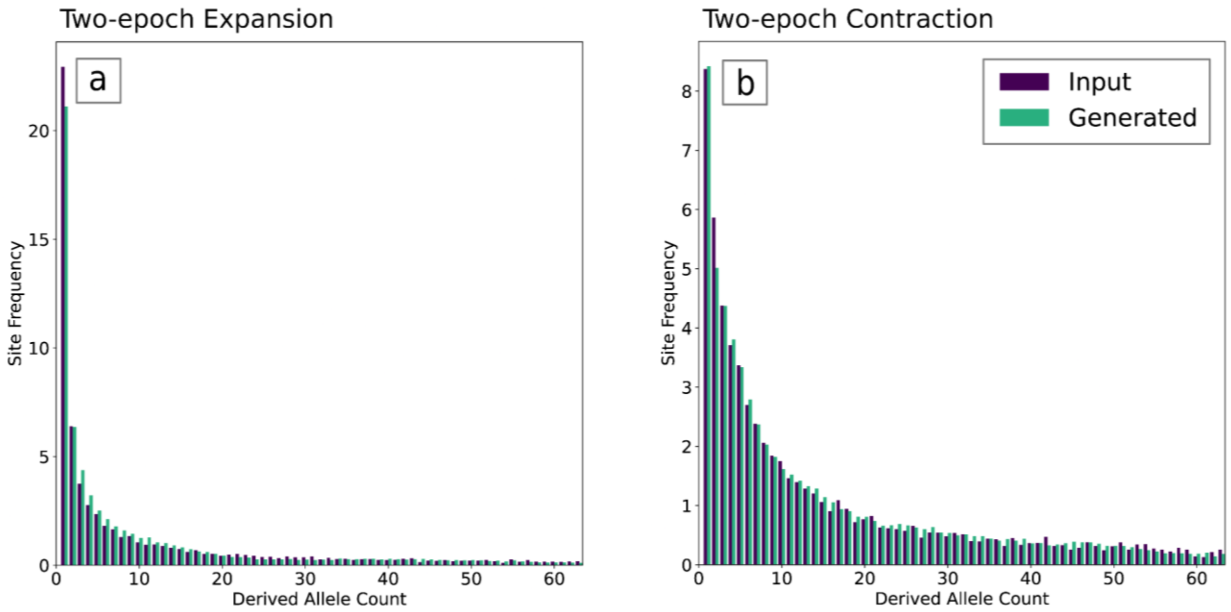


**Supplemental Figure 7.** Input and generated site frequency spectra (SFS) averaged across 1000 samples for generated (green) and input (purple) alignments under a (a) 10-fold population expansion and a (b) 10-fold population contraction.

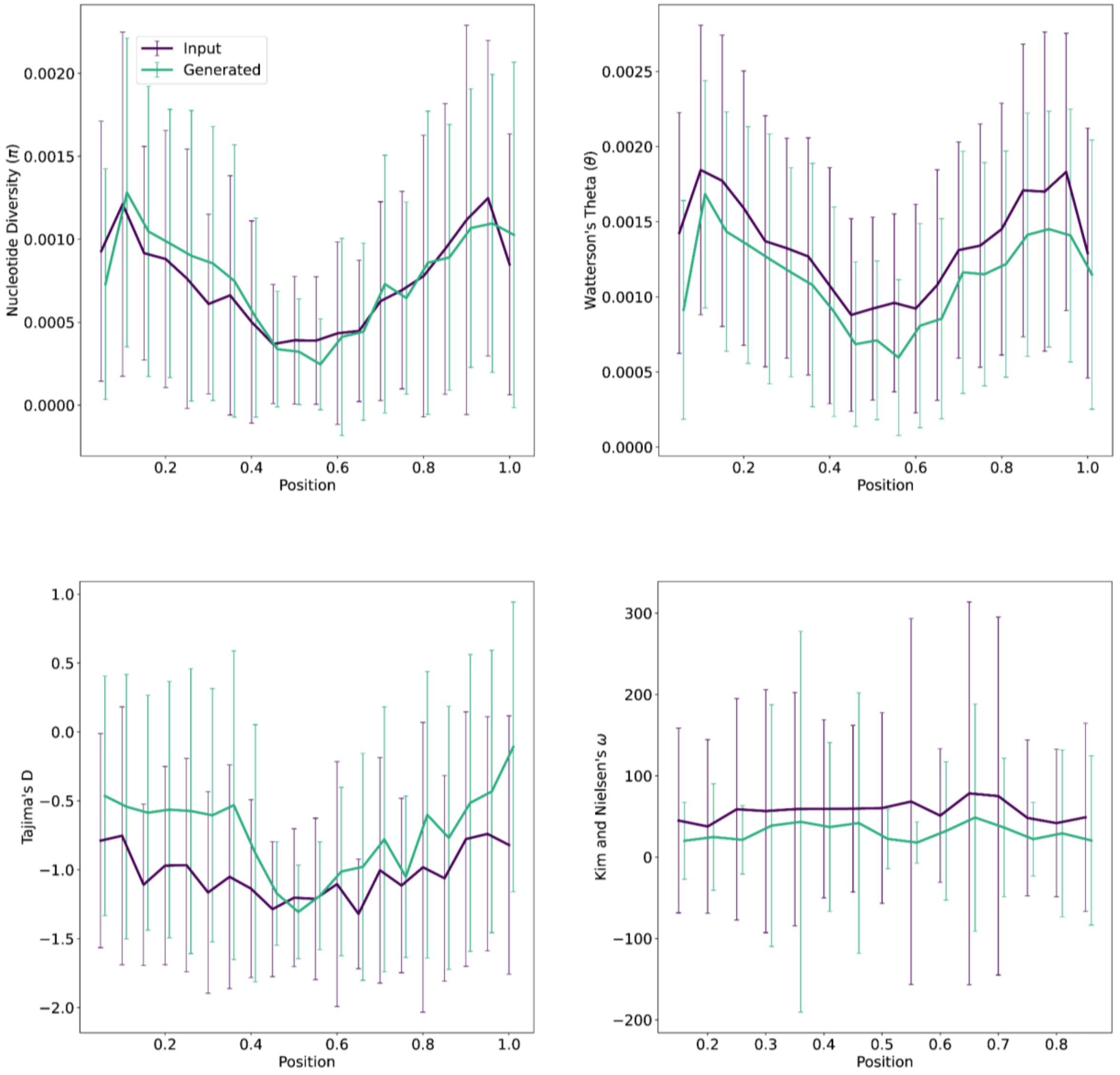


**Supplemental Figure 8.** Windowed diversity statistics for input (purple) and generated (green) alignments under a model of moderate selection. Error bars represent standard deviation. Nucleotide diversity (pi), Watterson's theta, Tajima's D are calculated in non-overlapping windows of 5% of the chromosome length. Kim and Nielsen's omega is calculated in overlapping windows 30% of the chromosome length and sliding 10% of the chromosome for each window.

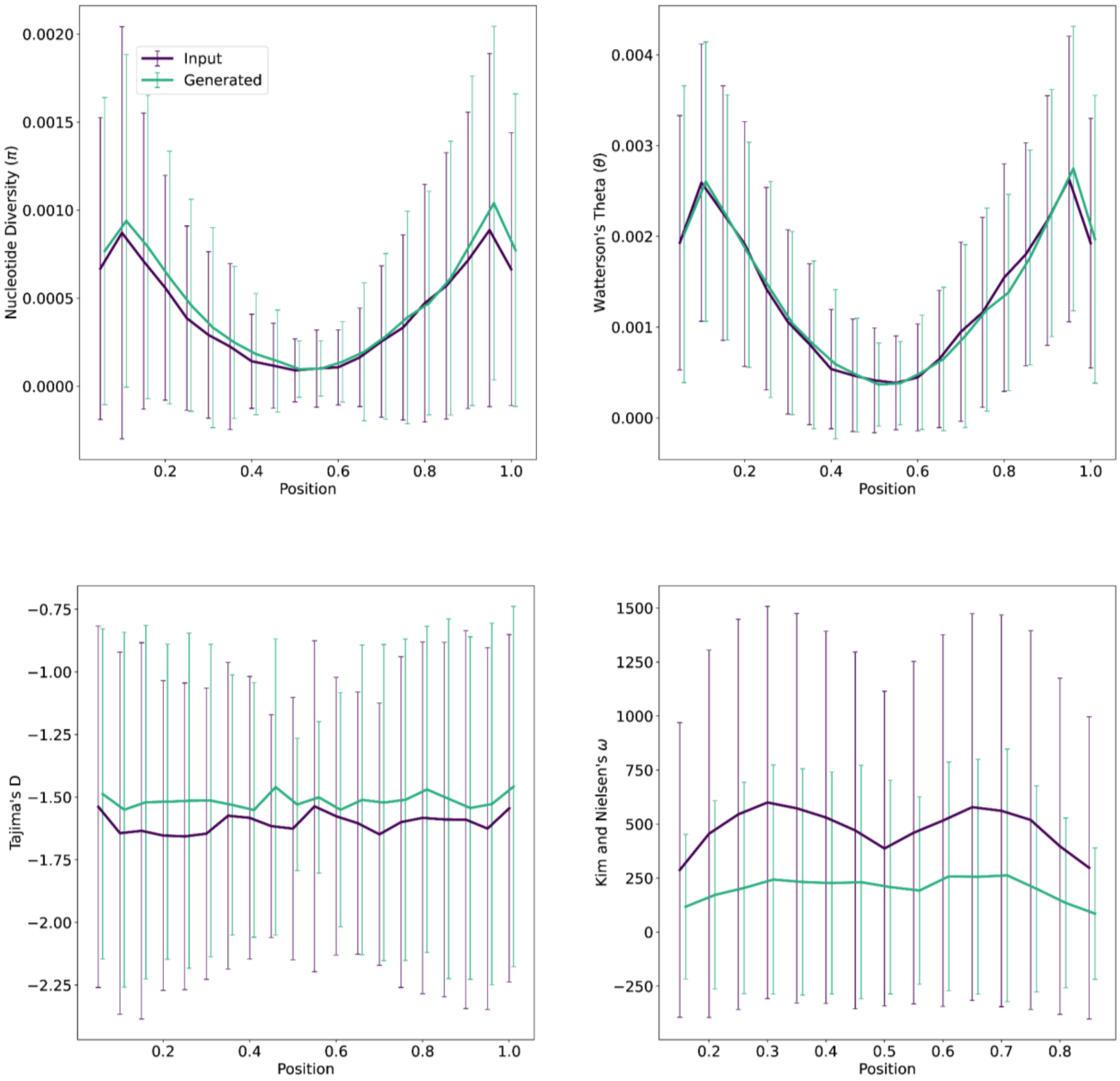


**Supplemental Figure 9.** Windowed diversity statistics for input (purple) and generated (green) alignments under a model of strong selection. Error bars represent standard deviation. Nucleotide diversity (pi), Watterson's theta, Tajima's D are calculated in non-overlapping windows of 5% of the chromosome length. Kim and Nielsen's omega is calculated in overlapping windows 30% of the chromosome length and sliding 10% of the chromosome for each window.

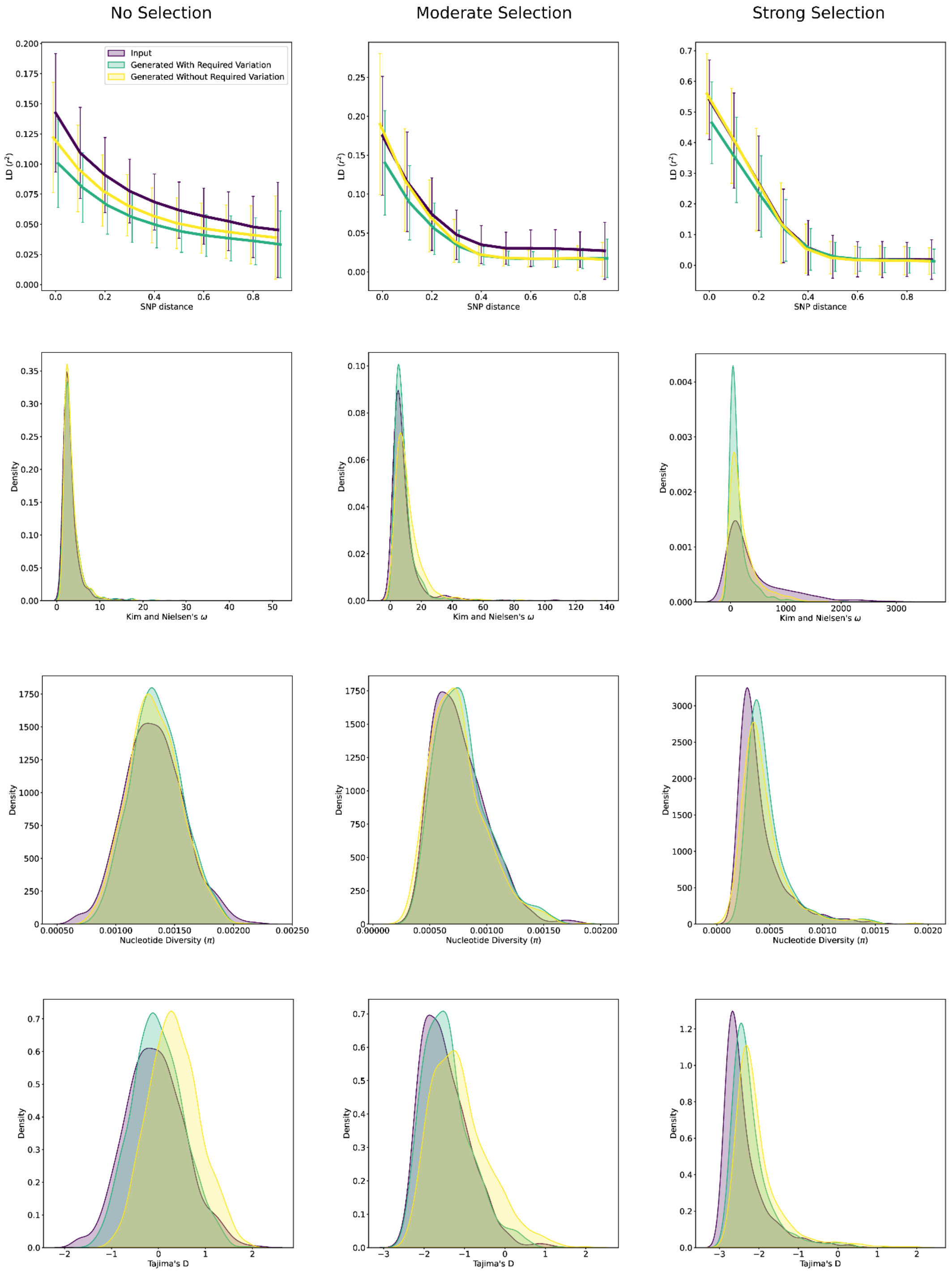


**Supplemental Figure 10.** GAN performance when invariant sites are dropped from the alignment compared (without required variation) to alignments where variation is forced through weighted random sampling (with required variation) for models of no selection (standard neutral with recombination), moderate selection, and strong selection. Each statistic shows the average over windows along the chromosome (LD) or distribution ($\omega$, $\pi$, Tajima's D) across 1000 alignments.

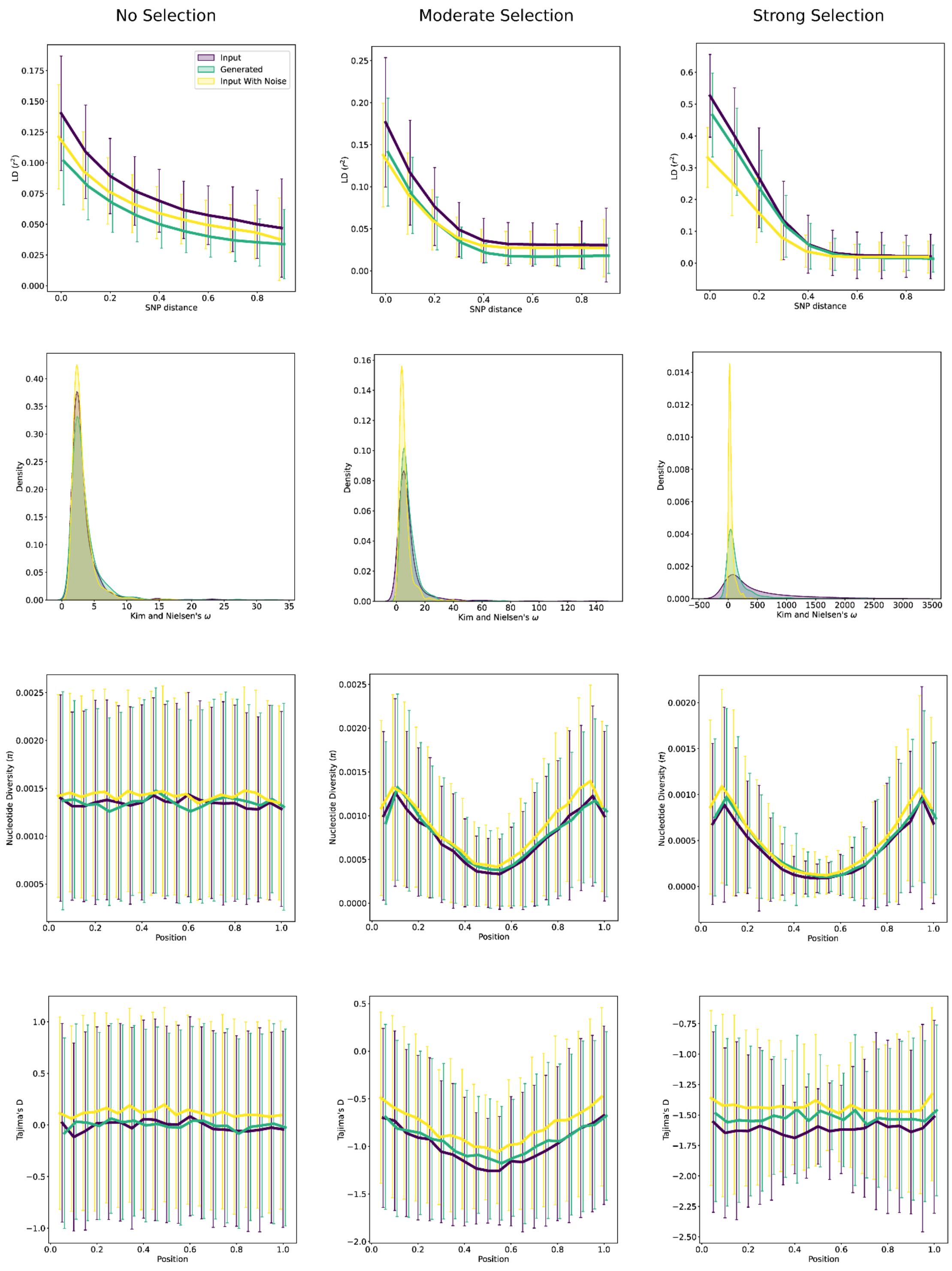


**Supplemental Figure 11.** Comparison of input alignments and input alignments with one percent of values flipped (input with noise) to GAN generated alignments for models of no selection (standard neutral with recombination), moderate selection, and strong selection. Each statistic shows the average over windows along the chromosome (LD, $\pi$, Tajima's D) or distribution ($\omega$) across 1000 alignments.

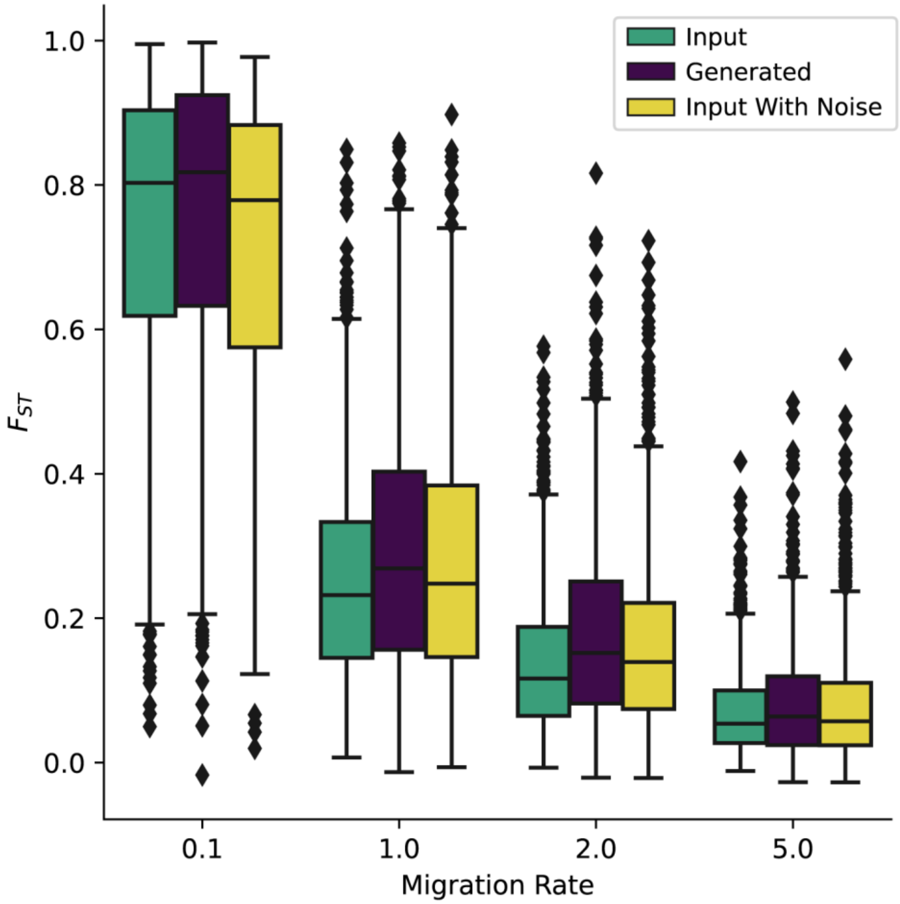


**Supplemental Figure 12.** Fixation index ($F_{ST}$) as a function of migration rate (shown in units of $4Nm$) in a comparison of input alignments and input alignments with one percent of values flipped (input with noise) to GAN generated alignments for subdivided population models. Each GAN was trained on alignments of two populations of 32 individuals and one of four bidirectional migration rates. Box and whisker plots of $F_{ST}$ were generated using 1000 alignments for each migration rate.
